# Long-read assembly of the Chinese rhesus macaque genome and identification of ape-specific structural variants

**DOI:** 10.1101/692749

**Authors:** Yaoxi He, Xin Luo, Bin Zhou, Ting Hu, Xiaoyu Meng, Peter A. Audano, Zev N. Kronenberg, Evan E. Eichler, Jie Jin, Yongbo Guo, Yanan Yang, Xuebin Qi, Bing Su

## Abstract

Rhesus macaque (*Macaca mulatta*) is a widely-studied nonhuman primate. Here we present a high-quality *de novo* genome assembly of the Chinese rhesus macaque (rheMacS) using long-read sequencing and multiplatform scaffolding approaches. Compared to the current Indian rhesus macaque reference genome (rheMac8), the rheMacS genome assembly improves sequence contiguity by 75-fold, closing 21,940 of the remaining assembly gaps (60.8 Mbp). To improve gene annotation, we generated more than two million full-length transcripts from ten different tissues by long-read RNA sequencing. We sequence resolve 53,916 structural variants (96% novel) and identify 17,000 ape-specific structural variants (ASSVs) based on comparison to the long-read assembly of ape genomes. We show that many ASSVs map within ChIP-seq predicted enhancer regions where apes and macaque show diverged enhancer activity and gene expression. We further characterize a set of candidate ASSVs that may contribute to ape- or great-ape-specific phenotypic traits, including taillessness, brain volume expansion, improved manual dexterity, and large body size. This improved rheMacS genome assembly serves as an ideal reference for future biomedical and evolutionary studies.

## Introduction

Rhesus macaque (*Macaca mulatta*) is the most widely studied nonhuman primate (NHP) in biomedical science (Gibbs, et al. 2007), and the closest human relative where gene editing has been approved for generating animal models. This makes it an indispensable species for understanding human disease and evolution, and as a result, a high-quality rhesus macaque reference genome is a prerequisite. The captive-born Chinese and Indian rhesus macaques represent two subspecies populations with moderate genetic differentiation (diverged about 162,000 years ago (Hernandez, et al. 2007)). Previous efforts using next-generation sequencing (NGS) have established a draft genome assembly of the Indian rhesus macaque (Gibbs, et al. 2007; Holt and Yandell 2011; Zimin, et al. 2014), but the current genome version (rheMac8) is incomplete with >47,000 gaps (∼72 mega-base pairs (Mbp) in length). Similarly, only a draft genome of the Chinese rhesus macaque was available (Yan, et al. 2011) (contig N50: 13 Kbp; total gap length: 331 Mbp). Due to the poor contiguity (fragmentation) and incompleteness (many gaps) of the current genomes, it has been difficult to systematically identify structural variants (SVs) in the macaque genome, which are known to be important in primate evolution and disease (Alkan, et al. 2011a; Alkan, et al. 2011c; Feuk, et al. 2006).

Long-read sequencing and multiplatform scaffoldings provide an opportunity for assembling high-quality genomes—in part due to long reads (>10 Kbp), more uniform sequence error distribution of long-read sequencing (such as PacBio), and the availability of multiple scaffolding technologies (such as Bionano and 10X Genomics) (Bend, et al. 2016; Chaisson, et al. 2015; Gordon, et al. 2016; Kronenberg, et al. 2018; Seo, et al. 2016). Compared to the short-read data from NGS, the long-read data are useful in resolving complex genomic regions, such as the highly repetitive and GC-rich regions. Long-read sequencing data in particular increases sensitivity for the identification and sequence resolution of SVs (Chakraborty, et al. 2018; Gordon, et al. 2016; Huddleston, et al. 2017; Kronenberg, et al. 2018).

The emergence of apes during evolution (Hominoidea: a branch of Old World tailless anthropoid primate native to Africa and Southeast Asia) required a series of evolutionary innovations, including taillessness (Williams and Russo 2015), large body size (Smith and Jungers 1997), increased brain volume/complexity (especially in great apes and humans) (Barton and Venditti 2014; MacLeod, et al. 2003; Rilling and Insel 1998; Rilling and Insel 1999), and improved manual dexterity (Berthelet and Chavaillon 1993; Napier 1993; Napier and Napier 1967). Apes are the sister group of Old World monkeys, together forming the catarrhine clade. Recently, using PacBio data, we reported the high-quality assembly of great ape genomes, including potentially adaptive human-specific SVs (Kronenberg, et al. 2018). A high-quality genome of an Old World monkey species (e.g., rhesus macaque), however, was lacking and is required as an outgroup to identify and characterize the functional genetic changes that occurred in the common ancestor of the ape lineage.

Here, we present the first high-quality Chinese rhesus macaque genome (rheMacS) *de novo* assembly using long-read sequencing and multiple scaffolding strategies and contrast it with the Indian macaque genome (rheMac8). We identify 53,916 SVs in rheMacS, and by comparative genomic approaches, we focus on 17,000 ape-specific structural variants (ASSVs), identifying potentially functional SVs that may contribute to the major phenotypic changes during ape evolution. The rheMacS assembly provides one of the most complete Old World monkey reference genomes to date and an important genetic resource to the biomedical community.

## Results

### De novo assembly of the Chinese rhesus macaque genome

We chose an adult male Chinese rhesus macaque (*Macaca mulatta*) and extracted high-molecular-weight DNA from peripheral blood. We first sequenced and assembled the genome using SMRT long-read sequencing (100-fold genome coverage; average subread length of 9.7 Kbp) and FALCON (Figure S1, Table S1,2 and Methods). We then scaffolded the contigs utilizing Bionano data (101-fold genome-coverage) and generated *de novo* optical genome map. We filled the remaining gaps in the assembled genome using PBJelly (English, et al. 2012) and corrected the errors of the PacBio long reads using Arrow (Chin, et al. 2013). To further improve the accuracy of the genome assembly, Illumina whole-genome shotgun (WGS) short-read data (50-fold genome-coverage) and Pilon (Walker, et al. 2014) were used to correct remaining errors in the sequence contigs (Table S2 and Figure S2). Finally, the Hi-C data (105-fold genome-coverage) was used to anchor scaffolds to chromosome models (Table S3 and Methods).

For the purpose of gene annotation, we extracted total RNA from 16 tissues, including large intestine, lung, epididymis, liver, testis, muscle, bladder, prefrontal cortex (PFC), cerebellum, skin, spleen, kidney, stomach, small intestine, heart, and pancreas (Methods). We performed full-length transcriptome sequencing (also called isoform sequencing, Iso-Seq) by pooling the RNA samples from the ten tissues (Table S1) and generating 100 Gbp of Iso-Seq data (Methods). To assess expression of the novel gene models, we also generated short-read RNA sequencing (RNA-seq) for each of the 16 tissues, producing 185 Gbp data (9.04∼13.92 Gbp for each tissue).

In summary, we have generated a *de novo* assembly of the Chinese rhesus macaque genome (rheMacS) that represents 2.95 Gbp of the chromosomes with contig N50 length of 8.19 Mbp and scaffolds N50 length of 13.64 Mbp (Table 1). Compared with the previous rhesus assembly (rheMac8), the rheMacS shows much less fragmentation (348,493 vs. 4,741 sequence contigs, >98% reduction in total contig numbers), improving sequence contiguity by 75-fold (contig N50) (Table 1, Figure 1 and Figure 2A) and scaffold N50 length by three-fold (Table 1 and Figure S3).

**Figure 1.**
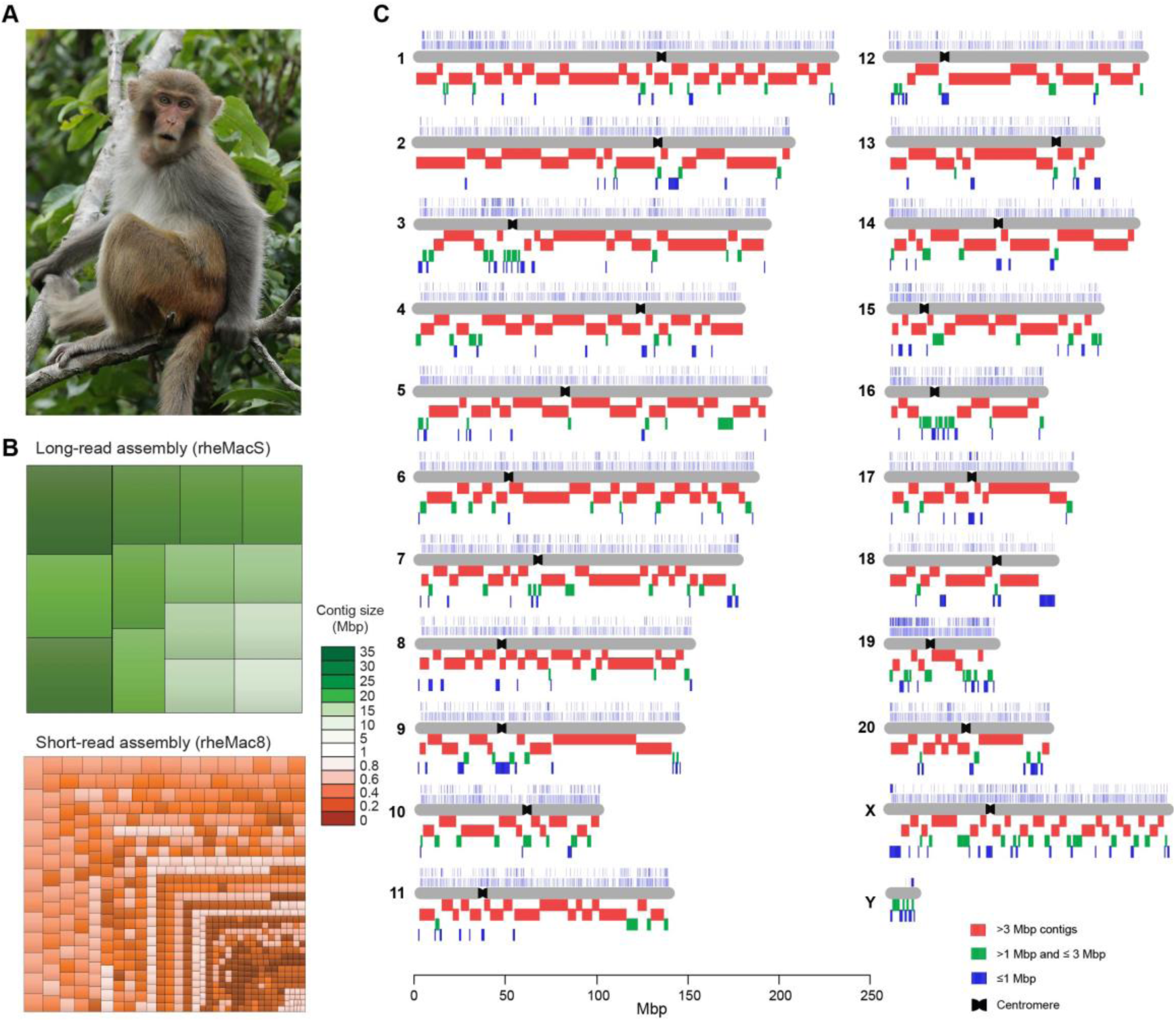
Long-read assembly of the Chinese rhesus macaque genome. **(A)** Chinese rhesus macaque (*Macaca mulatta*) (photograph courtesy by Jiaxin Zhao). **(B)** Treemaps for fragmentation difference between long-read and short-read rhesus assemblies. The rectangles represent the largest contigs that account for ∼300 Mbp (∼10%) of the assembly. **(C)** The chromosomal distribution of contigs of the rheMacS genome assembly. We compare genome sequence contiguity between rheMac8 and rheMacS. Thousands of gaps in rheMac8 were closed by longer contigs from rheMacS. The assembled contigs include >3 Mbp (red), between 1 Mbp and 3 Mbp (green), and those <1 Mbp (blue). The small contigs (<1 Mbp) tend to consist of either centromeric or telomeric sequences. The centromeres of each chromosome are indicated based on previous annotation (Rogers, et al. 2006).

**Figure 2.**
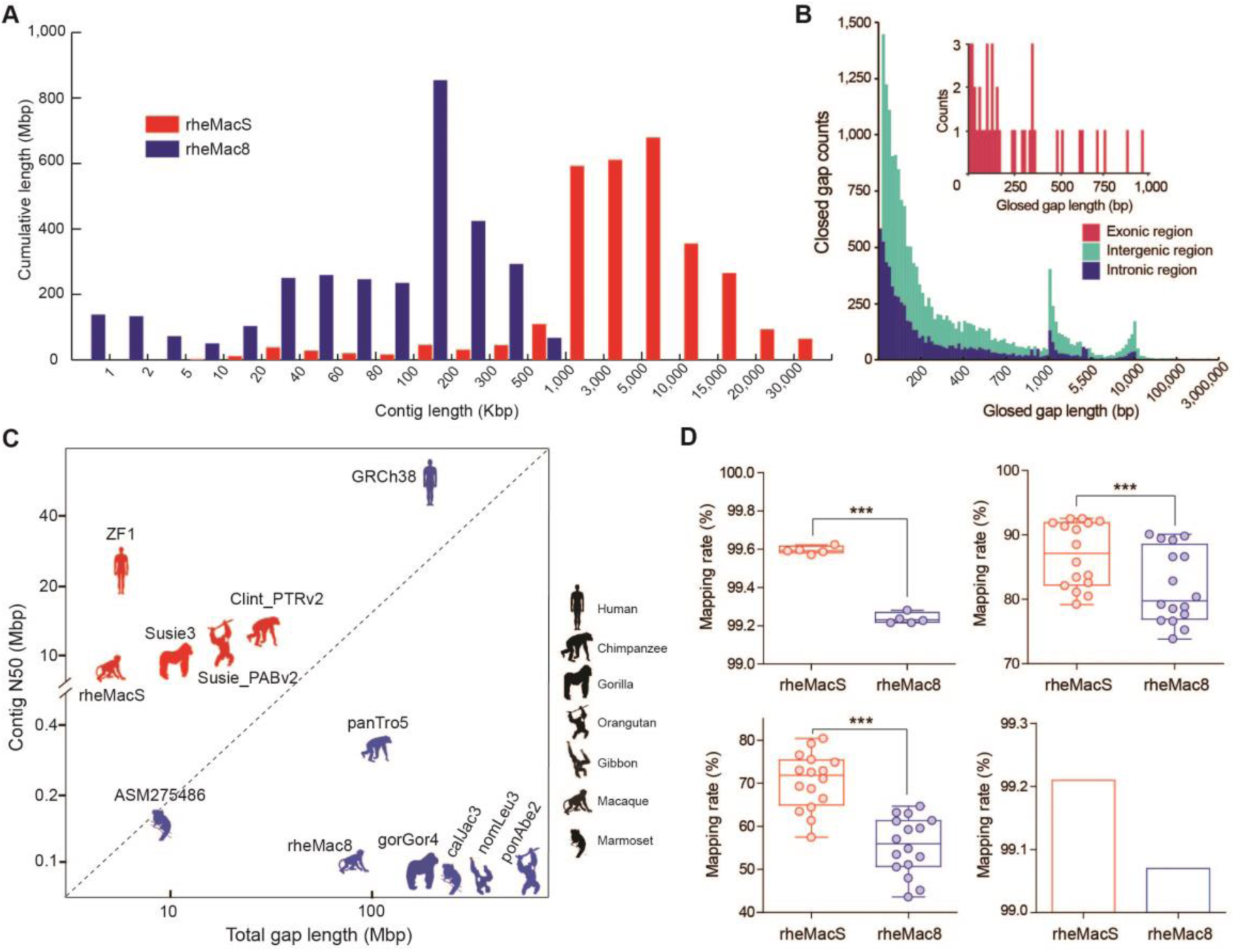
Quality assessment of the Chinese rhesus macaque genome assembly. **(A)** Comparison of sequence contig length distribution between rheMacS and rheMac8. **(B)** Length distribution of the closed gaps in rheMac8. **(C)** Comparison of assembly quality among various reported primate genome assemblies. Genomes assembled with long PacBio long reads (red) are compared against those assembled using Illumina short-read and Sanger sequencing data (blue) with the exception of common marmoset (ASM275486). **(D)** Comparison of mapping rates when short-read NGS data (upper left), RNA-seq data (by *Hisat2*: upper right; by *Bowtie2*: lower left), and Iso-Seq data (lower right) are mapped to rheMacS and rheMac8, respectively. Two-tailed paired t test was used for statistical assessment. ***-P<0.001.

**Table 1.** Comparison of assembly statistics between rheMac8 and rheMacS.

| Assembly | rheMac8 | rheMacS |
| --- | --- | --- |
| Assembly approach | WGS and BAC | WGS and Hi-C |
| Sequencing platform | Sanger, Illumina | PacBio, Bionano, Hi-C, Illumina |
| Number of contigs | 348,493 | 4,741 |
| Contig N50 (Mbp) | 0.11 | 8.19 |
| Number of scaffolds | 286,263 | 4,543 |
| Scaffold N50 (Mbp) | 4.19 | 13.64 |
| Number of gaps* | 47,882 | 2,858 |
| Total gap length (Mbp)* | 72.05 | 5.46 |
| Total bases (bp) † | 2,835,963,390 | 2,955,490,605 |
| Ungapped bases (bp) ‡ | 2,763,913,633 | 2,950,026,318 |

The assembly statistics of rheMac8 were obtained from NCBI (accession number: GCA\_000772875.3). WGS: whole-genome shotgun, BAC: bacterial artificial chromosome; \*only N-base regions in the assembled chromosomes were counted; †only chromosome-placed bases were counted; ‡non-N bases (bp) in the assembly.

### Quality assessment

We used the assembled rheMacS genome to first close gaps in the rheMac8 reference genome (Methods). We filled 21,940 of remaining N-gaps in rheMac8 adding 60.81 Mbp of additional sequences (2% of the entire genome, Table 1, Figure S4 and Table S4). As expected, 75% of the closed gaps consisted of various classes of repeat DNA: long interspersed nuclear elements (LINEs), short interspersed nuclear elements (SINEs), and simple short tandem repeats (Figure S5 and Table S6). Notably, 7,146 of the filled gaps (13.27 Mbp) map within genes and 41 map to the coding regions adding 10.17 Kbp of protein-encoding sequences (Figure 2B and Tables S4, S5).

The rheMacS assembly has far fewer gaps (rheMacS: 2,858 vs. rheMac8: 47,882), which are shorter in length (rheMacS: 5.46 Mbp vs. rheMac8: 72.05 Mbp) when compared to rheMac8 (Figure 2C and Table 1). To assess the consensus accuracy of rheMacS, we aligned each chromosome sequence of rheMacS to rheMac8. More than 98% of the assembled sequences are concordant by length and orientation (Figure S6 and Table S7). To evaluate sequence accuracy, we also mapped the 50-fold Illumina short-read data to the rheMacS assembly as the previous studies described (Bickhart, et al. 2017; Koren, et al. 2018). The estimated quality value (QV) score was 50 (1.11 × 10E-5 variants per base) (Table S8 and Methods). This is well below one error per 10,000 bases, a quality standard used for human genomes (Schmutz, et al. 2004).

Next, we compared the mappability between rheMacS and rheMac8 by deep sequencing genomes from five additional unrelated Chinese rhesus monkeys (Illumina X10 PE-150bp, 50× depth per individual). We mapped the short-read data to rheMacS and rheMac8, respectively, observing a higher mapping rate for rheMacS compared to rheMac8 (99.60% vs. 99.24%, respectively) (Figure 2D and Table S9). Genome-wide variant calling using the WGS data and different tools (GATK and SAMtools) for the five monkeys shows a similar number of SNV types (SNPs and INDELs) and SV types (deletion, insertion, duplication and inversion, SV length ≥50 bp) (Table S10). These results suggest that rheMacS will be useful for future variant detection.

Interestingly, when we mapped the RNA-seq short reads from the 16 tissues (∼37M reads per tissue) to rheMacS and rheMac8, we observed a significantly increased mapping rate for rheMacS compared to rheMac8 (86.87% vs. 81.96%, respectively) (Figure 2D and Table S11). Long-read Iso-Seq data show similar mapping rates (99.21% for rheMacS vs. 99.07% for rheMac8) (Figure 2D and Table S12). While some of these differences may relate to divergence between Chinese and Indian origin of the genomes, our results suggest that the rheMacS assembly will also enhance future rhesus monkey transcriptomic analyses.

### Gene Annotation

We performed *ab initio* gene annotation for rheMacS using the 100 Gbp of long-read (Iso-Seq) sequence data (2,468,473 full-length non-chimeric (FLNC) transcripts with mean length of 2,759 bp (Figure S7), and the 185 Gbp of short-read RNA-seq data generated from the 16 tissues (Methods). We annotated in total 20,389 protein-coding genes in rheMacS, just slightly fewer than rheMac8 (20,605) (Table S13). The analysis, however, generated more isoforms in rheMacS (288,773 in rheMacS vs. 276,000 in rheMac8) (Table S12) where the average length was longer for rheMacS (1,564 bp in rheMacS vs. 1,489 in rheMac8) (Figure S8 and Table S13). Improved gene annotation results, in part, from rescued missing exons and better coverage of full-length transcripts after gap closures (Table S5). Comparison of orthologous gene families between rheMacS and other primate genome assemblies (rheMac8, ape and mouse) indicated similar gene family composition (Figure S9), suggesting a reliable annotation quality for rheMacS. We also annotated repeats by RepeatMasker (Tempel 2012) and Tandem Repeats Finder (Benson 1999) (Methods). We found that 1.5 Gbp (54.04%) of the rheMacS genome were annotated as repeats, similar with the repeat ratios in rheMac8 and ape genomes (Tables S14, S15).

To further assess gene annotation, we benchmarked 4,104 universal single-copy orthologs (BUSCO) (Simao, et al. 2015), which by design should be uniformly present in all mammalian genomes. We find that 3,836 BUSCO genes (93.5%) are completely annotated in rheMacS. There are 191 BUSCO genes (4.7%) with fragmented annotation and only 77 BUSCO genes (1.8%) not annotated in rheMacS, contrasting the higher missing rate in rheMac8 (3.1%) (Table S16). We also annotate noncoding RNA (ncRNA) genes in rheMacS (Methods), obtaining 49,698 long noncoding RNA (lncRNA), 11,035 microRNA (miRNA), 544 ribosomal RNA (r|RNA), 2,373 small-nucleolar RNA (snRNA), and 718 transfer RNA (tRNA) (Table S17).

### Identification of SVs in the rheMacS genome assembly

Given the increased sensitivity for SV detection and their increased probability of affecting gene expression and phenotype (Kronenberg, et al. 2018; Sudmant, et al. 2015), we identified putative SVs (Sedlazeck, et al. 2018) by comparison against the previous Indian macaque genome assembly (rheMac8). We defined SVs as variants ≥50 bp in size and identified 53,916 SVs in the rheMacS assembly (Figure 3). The set includes 28,117 deletions, 24,431 insertions, 693 duplications and 675 inversions (Table S18, Methods). These rheMacS SVs correspond to 26% (1,399/5,456) of the previously reported SVs from Indian rhesus macaques using array genomic hybridization (aCGH) and Illumina short-read genome sequencing (Gokcumen, et al. 2013; Gordon, et al. 2016; Iskow, et al. 2012; Lee, et al. 2008). Notably, we estimated that 96% (51,919/53,916) of the SVs are novel, highlighting the advantage of long-read sequencing data for genome-wide SV detection (Figure 3E and Table S18).

**Figure 3.**
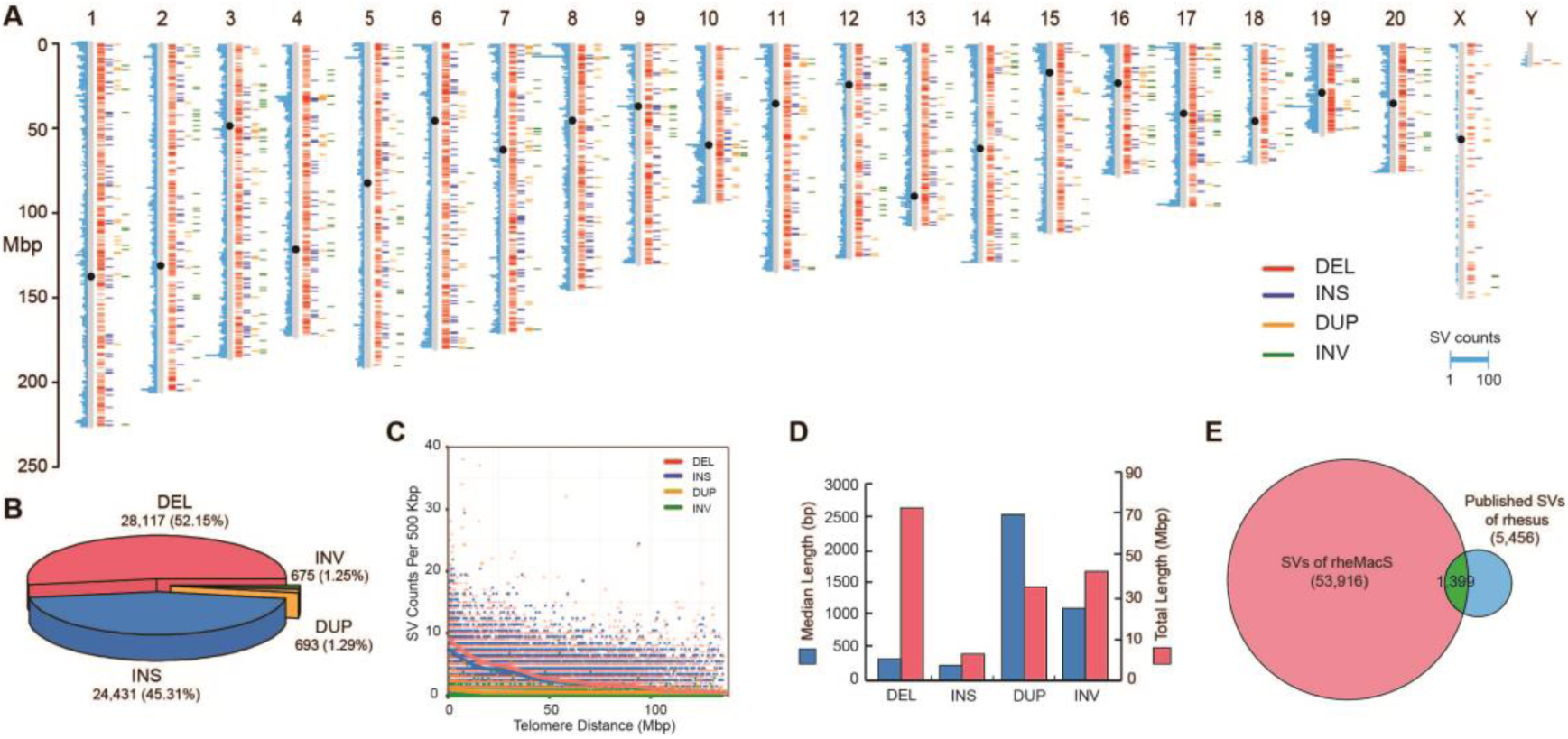
Structural variants (SVs) in rheMacS. **(A)** The distribution of large SVs (≥1 Kbp) among the rhesus macaque chromosomes. The histogram marks on each chromosome (light blue on the left) indicate the counts of SVs based on per 500 Kbp windows. The black dot on each chromosome indicates the centromere position. **(B)** The percentages of the four SV types including deletions (DEL), insertions (INS), duplications (DUP) and inversions (INV). **(C)** The SV distribution along with the increase of telomere distance. The SVs are counted with a sliding window size of 500 Kbp. The multi-color dots refer to the four-type SV counts in a 500 Kbp bin, and the solid lines indicate the distribution of average counts. **(D)** The length statistics of the rheMacS SVs. **(E)** Overlaps of the rheMacS SVs with previously reported SVs by aCGH and NGS data. The overlap cutoff is set to require >50% reciprocal overlapping of SV length.

The genomic distribution of the identified SVs in rheMacS is not random, and they tend to cluster in the proximal telomere regions (Figure 3A and Figure 3C), consistent with the pattern seen in our recent report of long-read human genomes (Audano, et al. 2019). We estimate a 1.8-fold increase in SV density within the subtelomeric regions (5 Mbp region of each chromosome end, p value < 10E-16, permutation). The median lengths for deletion, insertion, duplication and inversion are 319 bp, 231 bp, 2,575 bp and 1,119 bp, respectively. Combined, the SVs span 170.8 Mbp and affect 5.78% of the entire genome (Figure 3D). Using VEP (Variant Effect Predictor), we find that 2% (1,069 SVs) map within exons, 39% (21,009 SVs) in introns, and the remaining 59% (31,838 SVs) in intergenic regions. The newly identified >50,000 SVs by long-read sequencing data will be a useful resource in studying primate genome evolution.

To further validate these 53,916 SVs, we genotyped them in the five unrelated Chinese rhesus macaques with Illumina WGS data (Illumina X10 PE-150bp, 50× depth per individual) using SVTyper (Chiang, et al. 2015) (Methods). The result showed that 42,126 (78.13%) SVs could be validated, among which 5,654 (13.42%) were fixed and 36,472 (86.58%) were polymorphic (Figure S10 and Table S18). There were 11,790 (21.87%) invalidated SVs, and the majority of them (59.58%) were located in repeat regions, such as LINEs, SINEs and simple repeats, suggesting that short-read data is limited to resolve SVs in repetitive regions. In particular, 94.67% of the invalidated SVs were insertions, highlighting the limitation of NGS short-read data in detecting novel insertions (Alkan, et al. 2011b).

### Detection of ASSVs

Sequencing of both ape and macaque using the same long-read sequencing platform provides an opportunity to identify, for the first time, ASSVs that emerged in the ape lineage since divergence from the Old World monkeys. Using smartie-sv (Kronenberg, et al. 2018), we performed genome-pairwise comparisons among the rheMacS assembly and three published great ape long-read genome assemblies (chimpanzee, orangutan and gorilla) (Kronenberg, et al. 2018) as well as a human long-read assembly (ZF1) (Figure S11 and Table S19, Methods). To exclude SVs that occurred in the rhesus macaque lineage, we used the published genome assembly of common marmoset (assembly ID: ASM275486), which is the latest and most complete genome assembly of a New World monkey species (contig N50: 155 Kbp; total gap length: 9.5 Mbp) (Figure 2C). Considering the poor quality of the current gibbon genome assembly (contig N50: 35 Kbp; total gap length: 205 Mbp) (Figure 2C), we only used it when conducting local sequence alignment for manual check of the candidate ASSVs in order to distinguish ASSVs from great-ape-specific SVs (GASSVs).

Using the above strategy, we detected 17,000 candidate ASSVs after the filtering (Methods), including 13,456 deletions and 3,544 insertions. ASSVs account for only 7.97 Mbp (on average 0.26% of the ape genomes) (Figure 4 and Table S20). As expected, the majority of the ASSVs (78.05%) map to common repeats in the genome such as SINEs, LINEs and LTRs (Table S21). The ASSV chromosomal distribution is concordant with the known sequence synteny relationships between rhesus macaque and ape (Figure S12 and Table S22) (Rogers, et al. 2006).

**Figure 4.**
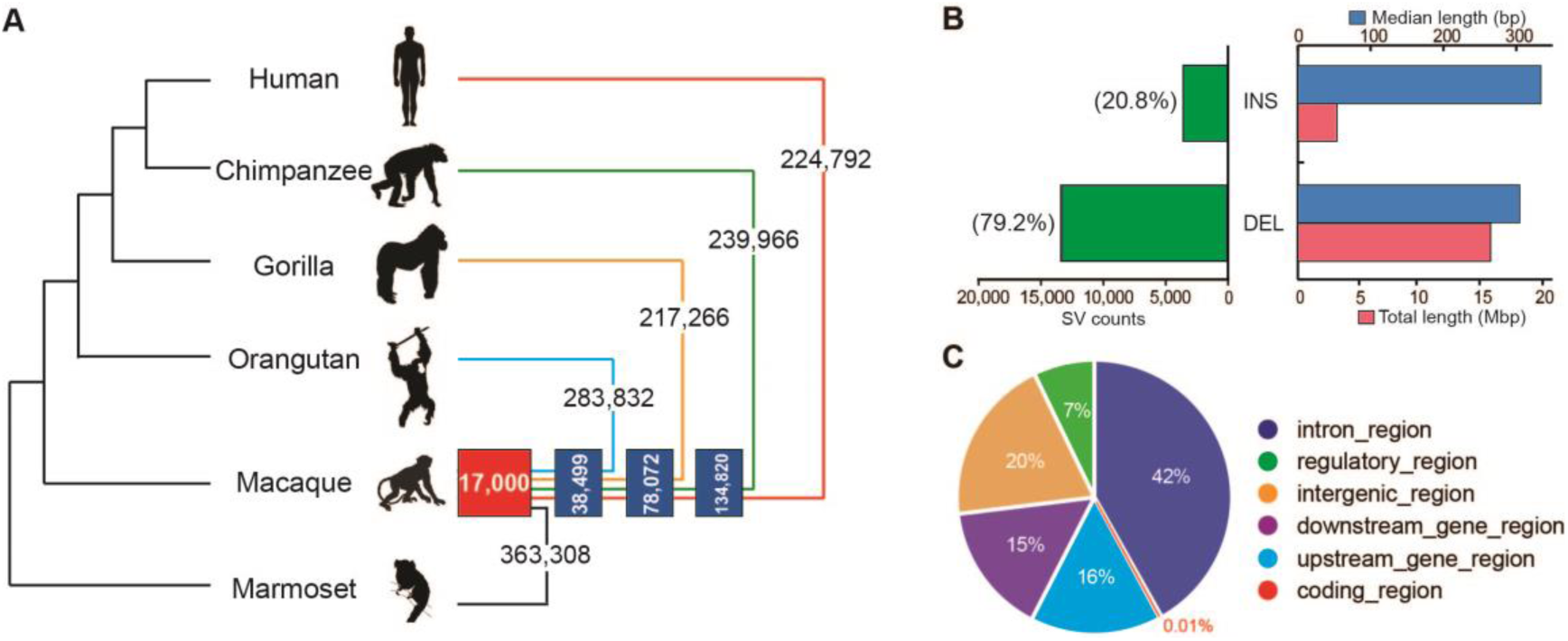
Summary of the ape-specific structural variants (ASSVs). **(A)** The cladogram depicts the phylogenetic relationship among the studied primate species, including human, three great apes, rhesus macaque, and common marmoset. The numbers of the identified SVs by genome-pairwise comparisons are indicated. The numbers in the blue boxes represent the overlap for different pairwise results. The number in the red box indicates the identified 17,000 ASSVs. Gibbon (lesser ape) was not included due to the poor quality of the published genome assembly. We used the common marmoset genome assembly generated by short-read sequencing data to filter the SVs that occurred in the rhesus monkey lineage (bottom panel). **(B)** Statistics of ASSVs. **(C)** Pie plot for ASSV annotation.

We first annotated the identified ASSVs using VEP according to the human GRCh38 coordinates (Methods). 12,255 ASSVs map either within or near (5 Kbp flanking transcriptional start and end) 3,412 coding genes with the remaining 4,745 ASSVs mapping to intergenic regions (Figure 4C). We explored functional enrichment among the 3,412 ASSV-related genes using DAVID GO (gene ortholog) analysis and pathway analysis. Although no functional category reached statistical significance, we observe several interesting functional categories among the top 5% (10/208) categories (Figure S13 and Table S23). Cilium assembly and morphogenesis ranked the top one category in the GO biological process and FAC (functional annotation clustering, enrichment score=3.13)—categories that play a critical role for development of the vertebrate nervous system and in regulating neuronal cell fate, migration and differentiation (Guemez-Gamboa, et al. 2014; Youn and Han 2018). We, therefore, speculate that ASSVs in this functional category might contribute to great ape brain-related phenotypic changes (e.g., expansion in brain size).

### ASSVs in gene-coding regions

There are 25 ASSVs (17 deletions and 8 insertions) located in the coding sequences of 32 genes (Table S24). We tested these 25 ASSVs by PCR and Sanger sequencing, and 16 of them were validated as true ASSVs and 9 were false ASSVs (Table S24). Among the 16 validated coding ASSVs (9 deletions and 7 insertions) (Table 2), 6 were annotated as NMD (nonsense-mediated decay) variants (Chang, et al. 2007), 4 as splice site variants, 2 as in-frame deletions, 2 as frameshift variants and 2 as coding variants. These SVs affect 15 genes, which are involved in brain function/neuro-diseases (*IL20RB*, *NMNAT3*, *CLCN3*, *FBF1*, *ZNF563*, *PPP1R15A* and *PTK6*), bone development (*VPS33A*, *CLCN3* and *DECR2*), spermatogenesis/sexual hormone (*EXOSC10* and *RUVBL2*), immunodeficiency (*UNG*), fatty-acid degradation (*DECR2*) and skin disease (*SLC17A9*) (Table 2). For example, there is an in-frame deletion ASSV (318 bp deletion) in *CCDC168*, a gene with unknown function (Figure S14 and Table 24). There are three ASSVs disrupting the splice acceptor/donor sites, resulting in lineage-specific protein (3,076 bp insertion in *NMNAT3*) (Figure S15 and Table S24) and lineage-specific transcript isoforms (316 bp insertion in *EXOSC10* and 1,435 bp insertion in *IL20RB*) (Figure S16 and Table S24). Functionally, *IL20RB* and *NMNAT3* are involved in neuroprotection (Abd Nikfarjam, et al. 2013; Kitaoka, et al. 2013). *EXOSC10* is associated with the regulation of male germ cells (Jamin, et al. 2017).

**Table 2.** The 16 validated ASSVs located in gene-coding regions.

| Chrom | Start | End | Length | Type | Gene | Exon | Consequence | Functions* |
| --- | --- | --- | --- | --- | --- | --- | --- | --- |
| chr1 | 11095424 | 11095738 | 316 | INS | *EXOSC10* | 4/5 | Splice_acceptor_variant | Spermatogenesis |
| chr3 | 136986715 | 136988148 | 1435 | INS | *IL20RB* | 3/7 | Splice_donor_variant | Neuroprotective and intelligence |
| chr3 | 139580804 | 139583878 | 3076 | INS | *NMNAT3* | 5/8 | Splice_acceptor_variant | Neuroprotective; intelligence and axonal protection |
| chr4 | 169663104 | 169663246 | 144 | INS | *CLCN3* | 2/14 | Coding_sequence_variant | Schizophrenia, autism spectrum disorder; bone pattern and synaptic transmission |
| chr12 | 109105086 | 109105279 | 195 | INS | *UNG* | 6/7 | NMD_transcript_variant | Immunodeficiency; epididymitis and somatic hypermutation |
| chr12 | 122264731 | 122264829 | 100 | INS | *VPS33A* | 2/14 | NMD_transcript_variant | Lysosome function; foot abnormality and acetabular dysplasia |
| chr13 | 102742592 | 102742592 | 318 | DEL | *CCDC168* | 4/4 | Inframe_deletion | <Unknown> |
| chr16 | 405554 | 405554 | 66 | DEL | *DECR2* | 3/10 | NMD_transcript_variant | Fatty-acid degradation; hemoglobin concentration and body mass index |
| chr17 | 75909754 | 75909754 | 304 | DEL | *FBF1* | 29/29 | NMD_transcript_variant | White matter hyperintensity; brain and eye measurements and cilium assembly |
| chr17 | 75909803 | 75909803 | 631 | DEL | *FBF1* | 29/29 | NMD_transcript_variant | White matter hyperintensity; brain and eye measurements and cilium assembly |
| chr19 | 12318606 | 12318606 | 317 | DEL | *ZNF563* | 4/4 | Frame_shift | Intelligence; schizophrenia and educational attainment |
| chr19 | 48874925 | 48874925 | 285 | DEL | *PPP1R15A* | 1/1 | Inframe_deletion | White matter disease and Alzheimer's disease |
| chr19 | 49004220 | 49004220 | 337 | DEL | *RUVBL2* | 4/15 | NMD_transcript_variant | Chorionic Gonadotropin level; follicle stimulating hormone level; cell cycle and mitotic |
| chr20 | 62965134 | 62965134 | 564 | DEL | *SLC17A9* | 9/13 | Splice_region_variant | Porokeratosis; cutaneous photosensitivity; papule and pruritus |
| chr20 | 63533689 | 63533689 | 537 | DEL | *PTK6* | 4/8 | Frame_shift | Dysgraphia; schizophrenia, autism spectrum disorder and hair shape |
| chr22 | 39964324 | 39964903 | 578 | INS | *Z82206.1* | 2/2 | Coding_sequence_variant | <Unknown> |

Chrom: Chromosome; INS: insertion; DEL: deletion; NMD: Nonsense mediated decay. \*The gene functions are collected from GeneCard database and literatures (De Nys, et al. 2001; Kondo, et al. 2017; Larrouture, et al. 2015; Moon and Parker 2018; Perez-Duran, et al. 2012; Riazanski, et al. 2011; Shin, et al. 2017; Siep, et al. 2004).

### ASSVs in regulatory elements functioning in the brain

Since the majority of the identified ASSVs map to intronic or intergenic regions, their potential functional impact may relate to gene expression regulation. Utilizing previously published brain ChIP-seq data from human, chimpanzee and rhesus macaque (Xu, et al. 2018), we searched for ASSVs mapping to enhancers where corresponding genes showed ape-specific distances, i.e., human and chimpanzee have similar enhancer activities while rhesus monkey shows significantly lower or higher activities (P<0.05, two-tailed unpaired t-test). We first identified 7,155 ape-monkey differential enhancers (ADEs) in eight brain regions (Table S26, Methods), where we observe significant differences of H3K27Ac signal (marker of enhancer activity) between apes (human and chimpanzee) and rhesus macaques in at least one brain region. When overlapping the 17,000 ASSVs with these ADEs, we identify 87 ADEs corresponding to 111 ASSVs, which have the potential to affect the regulation of 65 nearby genes (Figure S17 and Table S26).

Among the 111 ASSVs, 21 (14 deletions and 7 insertions) are high confident based on manual curation (Table S25), which affect 20 ADEs (Figure 5A). Interestingly, except for two ADE showing monkey-ape difference in more than one brain region, the other 18 ADEs show interspecies differences in only one brain region. With respect to enhancer activity, 13 ADEs are enhancer gains, where apes show significantly stronger enhancer signals than macaque, and the other 5 ADEs are enhancer losses (Figure 5A and Table S26). Notably, the ape-gained enhancers are dominant in 4 of the 5 brain regions except for PcGm (precentral gyrus), which contained 3 ADEs—all of which resulted in enhancer loss in the ape lineage. We observed the same pattern when including the 111 candidate ASSVs (Figure S17). Interestingly, we identify 10 ADEs (containing 10 ASSVs) near 10 genes, which are related with neuronal cell function, brain function and neurological diseases (Figure 5A and Table S26).

**Figure 5.**
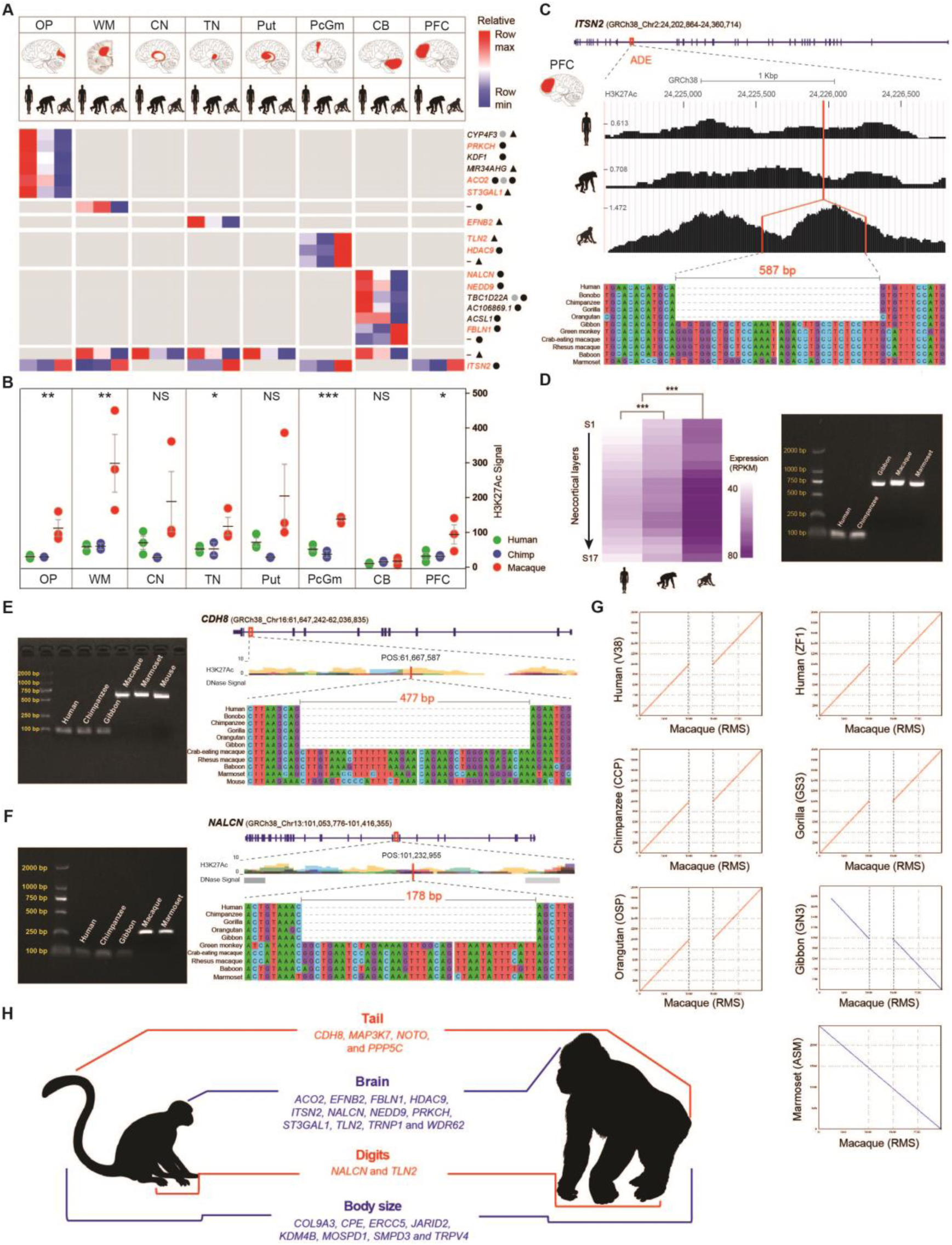
ASSVs associate with ape-specific (ASPs) or great-ape-specific phenotypes (GASPs). **(A)** Heatmap illustrating the ADEs with high-confident ASSVs in eight brain regions. The nearest genes are indicated and the corresponding brain regions are indicated in red. The neuro-function-related genes are highlighted in red. ASSV deletions (circles) and insertions (triangles) are denoted. The circles/triangles in black and gray refer to high-confident ASSVs and candidate ASSVs, respectively. **(B)** Comparison of H3K27Ac signals of the ADE with an ASSV (587 bp deletion) in *ITSN2* among human, chimpanzee and macaque. The ADE exhibits significant signal difference between human/chimpanzee and macaque in five brain regions (*-P<0.05; **-P<0.01; ***-P<0.001; NS-not significant, P>0.05). **(C)** A 587 bp deletion within intron-29 of *ITSN2* disrupts a putative enhancer sequence in the great ape lineage, with reduced enhancer activity in human and chimpanzee compared to rhesus macaque. The H3K27Ac signals in PFC and sequence alignments are shown. **(D)** *ITSN2* exhibits significantly lower expression in chimpanzee and human compared to macaque with 16 neocortical layers (1S-16S) and the adjacent white matter (17S) (left panel), consist with the enhancer difference in PFC. PCR validation is shown (right panel). **(E)** A 477 bp deletion located in intron-10 of *CDH8,* a gene related to tail development. **(F)** A 178 bp ape-lineage-specific deletion in intron-12 of *NALCN1*, a gene associated with human fetal adducted thumbs. **(G)** A dot plot alignment highlights a 587 bp deletion (located in *ITSN2*) in apes compared to rhesus macaque and marmoset. **(H)** Summary of candidate ASSVs/GASSVs located in genes associated with ASPs (in red) or GASPs (in blue). A statistical assessment of the H3K27Ac signals and gene expression difference was conducted using two-tailed unpaired and paired Student’s t-test, respectively. NS: not significant (P>0.05) and ∗∗∗: P < 0.0001.

We experimentally validated two ASSVs using PCR and Sanger sequencing. The 587 bp deletion that disrupts an ADE (GRCh38: chr2:24,224,548-24,226,881) in five brain regions of great apes in *ITSN2* (Figure 5B, Figure S18 and Table S26), which encodes Intersectin-2, affecting clathrin-mediated endocytosis and is critical in synaptic vesicle recycling in neurons (Pucharcos, et al. 2000). In the PFC, according to the published transcriptome data of 16 PFC cortical layers and the adjacent white matter (He, et al. 2017), the expression of *ITSN2* is significantly lower in humans and chimpanzees than in rhesus macaques, consistent with their reduced H3K27Ac signals (an indication of enhancer activity) due to the ASSV that disrupts an ADE in *ITSN2* (Figure 5C, 5D, 5G). Another 1,128 bp deletion is a cerebellum (CB)-specific ADE (GRCh38: chr6:11,262,523-11,265,271) of apes in *NEDD9* (neural precursor cell expressed, developmentally down-regulated 9), a gene relevant for dendritic spine maintenance, cognitive ability (Knutson, et al. 2016) and hindbrain development (Merrill, et al. 2004) (Figure S19 and Table S26). This ASSV leads to increased enhancer activities of *NEDD9* in cerebellum of humans and chimpanzees (Figure S19). Gene expression data of cerebellum in NHPs are not available to check its effect on *NEDD9* expression.

### ASSVs associated with ape-specific or great-ape-specific phenotypic traits

There are a set of ape-specific phenotypes (ASPs) that are not found among other primates, including taillessness (Williams and Russo 2015) and improved manual dexterity (Berthelet and Chavaillon 1993; Napier 1993; Napier and Napier 1967). In addition, brain capacity expansion (Barton and Venditti 2014; Rilling and Insel 1998) and large body size are great-ape-specific phenotypes (GASPs) (Hunt 2016; Rilling and Insel 1999; Smith and Jungers 1997). To understand whether the identified ASSVs might contribute to these ASPs or GASPs, based on their known functions from databases and published literature, we compiled a list of 451 genes related to these phenotypes (137 genes for tail development, 19 genes for brain size regulation, 56 genes for adducted thumbs and 239 genes for body size; Table S27 and Methods).

We performed a simulation to test the randomization of ASSVs for these ASPs or GASPs (see Methods for details). The results showed significance for body size (P=6.0E-06) and brain size (P=3.81E-04) but not manual dexterity (P=8.8E-01) or tail development (P=8.99E-01). This result suggests that there is an enrichment of ASSVs associated with GASPs.

To search for candidate ASSVs that potentially affect the 451 ASP/GASP-related genes, we conducted a detailed survey in each gene set of the four traits. Among the 137 tail-development-related genes (Table S27, Methods), we identified 32 ASSVs and 4 of them are high-confident ASSVs (Table S28). For example, we found a 477 bp deletion located in *CDH8* (Figure 5E). The mouse *Cdh8* knockout showed abnormal tail movements (Suzuki, et al. 2007). Another example is a 130 bp deletion in *MAP3K7* (Figure S20). The *MAP3K7* knockout mouse had a shorter tail. Importantly, the knockout mouse also showed an abnormal form of vertebral bodies (HP:0003312) and abnormal dental morphology (HP:0006482) (Zhao, et al. 2017), both of which show diverged phenotypes between monkeys and apes (Hunt 2016). In addition, we observed H3K27Ac peaks (human data) in these two ASSV regions (from ENCODE databases; Figures 5E and S20), implying that these ASSVs may contain regulatory elements such as enhancers. Notably, both genes belong to the Wnt-signaling pathway, a crucial pathway for tail extension and axial termination in vertebrates (Rashid, et al. 2014). Interestingly, the two genes are also highly expressed in the nervous system (brain and spinal cord) and are involved in body plan and somitogenesis during embryo development (Diez-Roux, et al. 2011).

The increase in ape manual dexterity involves a series of anatomical changes of the hand, including the evolution of flat nails and more sensitive finger pads, especially those associated with opposable thumbs (Berthelet and Chavaillon 1993; Marzke 1997; Napier 1993; Napier and Napier 1967). Among the 56-known adducted-thumb associated genes (ATA genes) (Human Phenotype Ontology ID: HP:0001181), we identify 12 ASSVs (6 deletions and 6 insertions) in 3 ATA genes of which 4 are high confident (Tables S27, S28). For example, an ASSV (a 178 bp deletion) that maps to the intron-12 of *NALCN* (sodium leak channel, non-selective) exhibited enhancer signals. It was reported that *NALCN* was associated with human fetal adducted thumbs (Bend, et al. 2016) (Figure 5F).

It is known that great apes have a larger body size compared to Old World monkeys and lesser apes (Hunt 2016; Smith and Jungers 1997). Among the 239 genes associated with body size (Table S27), we identified three genes (*CPE*, *COL9A3* and *ERCC5*) with GASSVs (Figures S21, S22). For example, *COL9A3* and *ERCC5* associate with a series of disorders showing growth failure and short stature (Jaarsma, et al. 2013; Paassilta, et al. 1999), and embryonic and postnatal growth retardation was previously reported in gene knockout mice (Harada, et al. 1999). These genes may affect body size in the great ape lineage as these GASSVs are shared among all great apes (Figures S21, S22, Tables S27, S28).

Taken together, we identify a set of ASSVs/GASSVs as candidate genetic changes associated with ape-specific or great-ape-specific traits that can serve as a resource to study the genetic basis of phenotypic innovations that emerged during ape–great ape evolution (Figure 5H). However, due to the lack of ENCODE data of the correspondent tissues in NHPs, the speculated regulatory roles of these ASSVs in the ape lineage are yet to be validated.

## Discussion

Rhesus macaque serves as an indispensable species for understanding human biology. Here, using long-read sequencing and multiple scaffolding technologies, we generated a high-quality genome assembly of rhesus macaque. The rheMacS genome assembly greatly improves the contiguity and completeness of the current version of the rhesus macaque reference genome (rheMac8). Using the rheMacS assembly, we characterized a large number of SVs, most of which were not observed in previous studies using array and NGS platforms. In particular, through comparative genomic analyses, we discovered 17,000 candidate ASSVs that may contribute to the emergence of ape-specific or great-ape-specific traits (such as taillessness, brain size, adducted thumbs and body size) during evolution (Figure 5H).

Among the 17,000 identified ASSVs, about 80% are deletions, and only 20% are insertions, indicating that losses of genomic sequences are more frequent than sequence gains, consistent with the results from the NGS data of great apes (Prado-Martinez, et al. 2013). Notably, 78.05% of the ASSVs contain repeat elements such as SINEs, LINEs and LTRs (Table S21), suggesting that repeat-element-mediated mutations are likely the main source of SVs. It should be noted that we only focused on the functional inference of two common SV types (deletions and insertions) because they account for the great majority (>98%) of SVs in the genome assemblies. Also, the ASSV filtering using NGS-based genome data of gibbon and marmoset can introduce unambiguity when calling the other SVs types such as duplications and translocations. These SVs types (duplications, inversions, translocations and chromosomal rearrangements) may also contribute to phenotypic evolution in primates, and they are worth studying in the future.

We focused on understanding the functional implications of ASSVs disrupting gene-coding integrity and those located in regulatory elements, which may contribute to ape-specific or great-ape-specific traits. We validated 16 ASSVs located in the coding regions of 15 genes, and 7 of them are associated brain functions and neurologic diseases (*IL20RB*, *NMNAT3*, *CLCN3*, *FBF1*, *ZNF563*, *PPP1R15A* and *PTK6*). For the ASSVs in the noncoding regions, we evaluated their potential functional effects based on the published brain ChIP-seq and transcriptome data. For example, we identified a 587 bp great-ape-specific deletion in an ADE of *ITSN2*, which may explain the attenuated enhancer activity in human/chimpanzee compared to macaque in five brain regions (OP, WM, TN, PcGm and PFC) (Figure 5B). Consistently, human and chimpanzee exhibited lower expression than macaque in the PFC and the adjacent white matter (Figure 5D, S18). Previous studies showed that overexpression of *ITSN2* resulted in the inhibition of transferrin uptake and the blockage of clathrin-mediated endocytosis, which is critical in synaptic vesicle recycling in neurons (Pucharcos, et al. 2000) and dendritic spine development (Nishimura, et al. 2006). Interestingly, *ITSN2* knockout mice displayed decreased grip strength and vertical activity (MGI:5631233), a trait that showed difference between apes and monkeys (Hunt 2016).

Primate evolution is an intricate process, and the changes of anatomical features from monkeys to apes may interact with each other during ontogeny and scale properly with body size as the organism grows. For example, most of the skeletal traits correlate strongly and positively with body size (Hunt 2016). In particular, the tailless trait of apes strongly linked with body size, positional behavior and bone structure, etc. (Fleagle 2013). Although it is hard to tell the order of trait emergence, it is possible that the related traits may be controlled by linked genetic changes during evolution. For example, the 178 bp deletion in *NALCN* (a gene related with adductive thumbs) disrupted an ADE in the cerebellum and exhibited a sharp difference in enhancer activities between monkeys and apes (Figure 5F and Table S26). It was reported that *NALCN* was also associated with neurodevelopmental diseases (Bramswig, et al. 2018) and the mutation of *NALCN* leads to syndromic neurodevelopmental impairment (Fukai, et al. 2016). Another example is the ASSV (242 bp deletion) within an ADE in PcGm in *TLN2* (Figure 5A), a brain-function-related gene (Gusareva, et al. 2018; Mendez-David, et al. 2017). This gene was also reported as a cause of fifth finger Camptodactyly (Deng, et al. 2016). Currently, the ENCODE data in NHPs are mostly from the brain, while the data from other tissues are not available. Given regulatory elements such as enhancers usually function in a tissue-specific manner, there is a great need to generate ENCODE data of NHPs so that the potential functions of the identified ASSVs associated with the key ape-specific traits other than the brain (such as taillessness and body size) can be annotated.

We used the marmoset assembly to filter out the monkey-lineage-specific SVs. Because of the poor assembly quality of marmoset, we found that 54.51% (9,266/17,000) ASSVs are uncertain (Table S20 and Methods). We conducted a manual check of 311 ASSVs, among which 16.4% (51 ASSVs) were high-confident ASSVs, and 18.3% (57 ASSVs) were false ASSVs. The remaining 65.3% (203 ASSVs) ASSVs were uncertain because the assembly and the sequence quality around the genomic regions of these ASSVs were poor in marmoset (Tables S24, S26, and S28). Similarly, when using the gibbon assembly to distinguish the GASSVs from ASSVs, we encountered the same situation. In contrast, we experimentally tested the 25 coding ASSVs, and 16 (64%) of them were validated by PCR (Table 24). The higher rate of high-confident coding ASSVs is likely due to the relatively conserved and well-resolved coding sequences in the genome assemblies. Nevertheless, the high-quality genome assemblies of New World monkeys (e.g. marmoset) and gibbons are needed to capture all lineage-specific SVs in apes.

High-quality genome assemblies are a prerequisite for comparisons among species. Most of the current primate genome assemblies were based on NGS short reads or shotgun Sanger sequencing with limited resolution for SV detection. For example, we observed a significantly poor reciprocal validation when comparing the NGS genome to the long-read assembled genome of rhesus macaque (Figure S23). Consequently, we should upgrade the current primate genome assemblies using long-read sequencing and multiple scaffolding techniques, a critical step forward in order to heighten the utility of NHP models for biomedical research and to greatly promote the understanding of primate evolution.

## Methods

### Sample Information

An adult Chinese rhesus macaque (*Macaca mulatta*) (male, five-year-old) was used for tissue sample collection in this study.

### Data generation

For long-read sequencing data, we extracted the high-quality genomic DNA from fresh blood samples of the macaque. The libraries with an average insert size of 20 Kbp were constructed using SMRTbell^TM^ Template Prep Kits and then sequenced on a PacBio Sequel instrument at the Genome Center of NextOmics Bioscience Co., Ltd (Wuhan, China). A total of 53 SMRT cells were run on the PacBio Sequel system and 299.6 Gbp (subreads) of data (with 100-fold coverage of genome) for rheMacS were generated. The average length and the N50 length of long subreads are 9.7 Kbp and 14.7 Kbp, respectively.

For Bionano optical mapping, the Bionano Genomics (BNG) instrument *Saphyr* was used to generate optical molecules using restriction enzyme Nt.BspQ1. High-molecular-weight DNA (>200 Kbp) was extracted from whole blood, and we constructed a high-quality sequencing library according to the recommended protocols. A total of 304 Gbp of clean data was generated with, on average, 8.6 labels per 100 Kbp, and the molecular reads N50 length was 216.4 Kbp (Figure S24).

For Hi-C data, two Hi-C libraries were prepared as described previously (Lieberman-Aiden, et al. 2009). Libraries were sequenced on an Illumina X10, which generated 308 Gbp (PE150) data with 103-fold genomic coverage.

For long-read RNA sequencing (Iso-Seq), total RNA was extracted from 10 tissues (heart, liver, spleen, lung, kidney, muscle, brain, epencephala, testicle and stomach) using a TRIzol extraction reagent (ThermoFisher), subjected to the Iso-Seq protocol with different ratios of three library sizes (1-6 Kbp,1-4 Kbp and 4-6 Kbp), and sequenced by the PacBio Sequel sequencer.

For short-read DNA sequencing, DNA samples were subjected to TruSeq DNA library kit (Illumina) and sequenced on the Illumina Sequencer X10 with paired-end 150 bp sequence reads. In total, 162 Gbp of data were generated.

For short-read RNA sequencing, RNA samples were subject to the TruSeq mRNA library kit (Illumina) and sequenced on the Illumina HiSeq X sequencer. 185 Gbp data was produced in total. Total RNA was extracted from 16 tissues (large intestine, lung, epididymis, liver, testis, muscle, bladder, PFC, skin, spleen, kidney, stomach, small intestine, cerebellum, heart and pancreas) using a TRIzol extraction reagent (ThermoFisher).

### De novo genome assembly

We estimated genome size by k-mer distribution analysis with the program Jellyfish (v1.1.11) (Marcais and Kingsford 2011) (k = 17) by the Illumina short reads. The 17-mer curve exhibits a significant heterozygosity peak and repeat peak for rheMacS (Figure S25), and the estimated genome size of rheMacS was around 3.07 Gbp, and the heterozygosity of the genome was about 0.5% (Figure S25).

FALCON (v1.8.7) (Eid, et al. 2009) was used for *de novo* assembly with PacBio long reads by the following steps: (i) raw subreads overlapping for error correction; (ii) preassembly and error correction; (iii) overlapping detection of the error-corrected reads; (iv) overlap filtering; (v) constructing a graph from overlaps; and (vi) constructing contigs from graph. After error correction, where a length cutoff of 14 Kbp was used for initial seed reads mapping, and then the error-corrected reads were used to construct the assembly graph and to generate a draft assembly with a genome size of 3 Gbp with a N50 length of 4.75 Mbp.

Hybrid genome assembly was performed by the Bionano Saphyr software with the manufacturer-recommended parameters. We constructed an optical map by running the hybrid scaffold pipeline (Bionano Solve 3.1). Subsequently, we filled the gaps based on the scaffold level genome. All PacBio long reads were aligned to the scaffold genome by PBJelly software (English, et al. 2012).

To further improve the accuracy of the genome assembly, a two-step polishing strategy was used for the initial assembly. The initial polishing was performed with Arrow (Chin, et al. 2013) using PacBio long reads only. Because of the high error rate of PacBio raw reads, we also used Pilon (v1.20) (Walker, et al. 2014) to further improve the PacBio-corrected assembly with the highly accurate Illumina short reads.

Scaffolds within each chromosomal linkage group were then assigned using the Hi-C-based proximity-guided assembly. The original cross-linked long-distance physical interactions were then processed into paired-end sequencing libraries (Figure S26). First, all the reads from the Hi-C libraries were filtered by the HiC-Pro software (v2.8.1) (Servant, et al. 2015), and then the paired-end reads were uniquely mapped onto the draft assembly scaffolds, which were grouped into 22 chromosome clusters and scaffolded using LACHESIS software (Burton, et al. 2013) (Table S29).

### Gap closure

We closed the gaps in the reference genome of rhesus monkey (rheMac8) using the approach of a previous study (Bend, et al. 2016). A region consisting of continuous runs of Ns (N>1) in the rheMac8 chromosomes was defined as a gap. We extracted 47,882 rheMac8 gaps (Table S4) based on the BED file format, and the 5 Kbp of flanking sequences upstream and downstream of the gaps were mapped to the assembly by MUMmer (Kurtz, et al. 2004) (*nucmer -f -r -l 15 -c 25*). We defined a gap as closed when the following two criteria were met: (i) the two flanking sequences of a gap in rheMac8 could be both aligned to the rheMacS assembly and the aligned length of the flanking sequences is >2.5 Kbp and (ii) when recording the rightmost coordinates (RC) and leftmost coordinates (LC) of rheMacS, non-N bases were observed between LC and RC in rheMacS. The total bases between LC and RC were counted as the closed-gap length.

We evaluated the repeat content of filled gaps, which were mapped to whole-genome repeats (annotated by rheMacS). An effective overlap was defined with overlapping length reaching at least 50% of the reciprocal similarities (Table S6). In addition, we annotated the filled gaps by mapping to the annotation of rheMacS, including exonic, intronic, and intergenic regions (Tables S4 and S5).

### Assembly quality assessment

Consensus quality: to evaluate the concordance between rheMacS and rheMac8, MUMmer was used to perform pairwise alignment and calculate the sequence identity (*nucmer --mum -c 1000 -l 100*).

Sequence quality: we mapped all 50× depth Illumina short reads to the rheMacS assembly using the BWA-MEM module (Eid, et al. 2009). Then we used Picard to mask the PCR duplicates and generated the dedup.bam file. Variants were called by the Genome Analysis Toolkit (GATK, v3.6) HaplotypeCaller module. The SNPs and INDELs were filtered using the GATK VariantFiltration module with the following criteria, respectively: SNPs filtering: “QUAL <50; QD < 2.0; FS > 60.0; MQ < 30.0; MQRankSum < −12.5; ReadPosRankSum < −8.0; DP < 15”; INDELs filtering: “QUAL <50; QD < 2.0; FS > 200.0; ReadPosRankSum < −20.0; DP < 15”. As previous studies described (Bickhart, et al. 2017; Koren, et al. 2018), we used QV (quality value) score to assess the sequence quality, which represents a per-base estimate of accuracy and is calculated as QV = −10log10(P) where P is the probability of error. We counted the total number of the homozygous SNVs (SNPs+INDELs), which represent the sites with base error in the rheMacS assembly. The base-error rate was calculated as the number of homozygous sites divided by the total size of the rheMacS assembly (Table S8).

Mappability: 50× depth WGS data was generated by Illumina sequencer X10 (PE-150) for five rhesus monkeys, which was used to evaluate the mappability and availability for variant calling. We used FastQC and GATK to perform quality control (QC) for the Illumina WGS reads, and then we mapped these short reads to rheMacS and rheMac8 assemblies respectively using the BWA-MEM module. SAMtools and GATK standard pipeline were used to call variants. Delly (v0.7.7) (Rausch, et al. 2012) was used to call SVs. All underlying algorithm, tool versions, and command lines for variant calling and filtering are listed in Table S30.

### Gene annotation

For gene annotation, 119.01 million RNA-seq reads and 1,275,860 Iso-Seq reads were generated with a mean length of 2,791 bp (Figure S27).

Gene models were constructed with MAKER (v2.31.8) (Holt and Yandell 2011), which incorporated the *ab initio* prediction, the homology-based prediction, and the RNA-seq assisted gene prediction. For the *ab initio* gene prediction, the repeat regions of the rheMacS genome assembly were first masked based on the result of repeat annotation, and then SNAP (Korf 2004) and Augustus (v3.2.2) (Stanke, et al. 2006) were trained for model parameters from homolog genes of BUSCOs, and they were then employed to generate gene structures. BUSCO (v3.0.1) was used to conduct the mammalian BUSCO analysis (Simao, et al. 2015) with the Mammalia odb9 set of 4,104 genes. BUSCO was run against all complete protein-coding sequence sets of annotation in rheMacS.

For ncRNA annotation, the published RNA database Rfam (http://rfam.xfam.org/) was used to predict rRNA, snRNA and miRNA by sequence alignments. The tRNAscan-SE and RNAmmer tools were used to annotate tRNA and rRNA, respectively. LncRNAs were predicted using ncrna_pipeline (https://bitbucket.org/arrigonialberto/lncrnas-pipeline) (Figure S28).

### Transcript analysis

After obtaining the raw Iso-Seq data from the PacBio Sequel system, we performed data QC and identified the full-length reads (by cDNA primer detection and removal: existence and completeness of 5’ primer, 3’ prime and poly-A). After this procedure, we obtained 2,468,473 FLNC reads. The size distribution of the FLNC reads, which correspond to candidate full-length transcripts, are displayed in Figure S7. To further improve the accuracy of isoform sequence, we made clustering for the FLNC reads using ICE (iterative clustering algorithm) and obtained the unpolished consensus reads. Then both these unpolished consensus reads and non-full-length reads were finally polished using Arrow (Chin, et al. 2013). Based on the Arrow consensus predicted accuracy, only sequences with accuracy of more than 99% were deemed high quality. The statistics of consensus reads are shown in Table S31. To further improve the base accuracy of consensus reads, we made error correction with short-read RNA-seq data using LoRDEC (Salmela and Rivals 2014). The number of NGS-corrected reads is listed in Table S31.

To compare the transcripts of rheMacS, rheMac8 and human, all Iso-Seq FLNC reads and ICE transcript models were aligned to the assembled genomes of rheMacS, rheMac8 and human (GRCh38) using GMAP (Wu and Watanabe 2005). The number of mean consensus number per isoform and the collapse isoforms were counted after filtering by identity and coverage (default arguments: identity rate > 0.99 and coverage rate > 0.95). In addition, to compare the annotation quality between rheMac8 and rheMacS, the RNA-seq short reads from 16 tissues were mapped to rheMac8 and rheMacS using Bowtie2 (v2.3.4.3) (Langmead and Salzberg 2012) and HISAT2 (Kim, et al. 2015), respectively (Table S11).

### Repeat analysis

Repetitive sequences including tandem repeats and transposable elements (TEs) were identified. First, we used Tandem Repeats Finder (TRF, v4.07b) (Benson 1999) to annotate the tandem repeats with options “*5 5 5 80 40 20 10 -m -ngs -h*”. Then, TEs were identified using RepeatMasker (v4.0.6) (Tempel 2012) to search against the known Repbase TE library (Repbase21.08) (Kapitonov and Jurka 2008).

### rheMacS SV detection

For long-read PacBio data, we used the mapping software NGLMR (Sedlazeck, et al. 2018) to align the subreads to the rhesus macaque reference genome rheMac8. Then Sniffles (Sedlazeck, et al. 2018) was used to call SVs from the bam file. For the NGS short-read Illumina data, we mapped the reads to rheMac8 by using BWA (Eid, et al. 2009). After sorting and removing duplicates, we used Delly (Rausch, et al. 2012) to call SVs. IGV (Integrative Genomics Viewer, http://software.broadinstitute.org/software/igv) was used for SV visualization.

The array- and NGS-based SV sets of rhesus macaque were obtained from dbVar (https://www.ncbi.nlm.nih.gov/dbvar) and previous studies (Gokcumen, et al. 2011; Gokcumen, et al. 2013; Iskow, et al. 2012; Lee, et al. 2008). The SVs were lifted over to the rheMac8 coordinates using LiftOver tool, and only the lift-over SVs were included in downstream analyses. The same SV was defined with overlapping length reaching at least 50% of the reciprocal similarities between published SVs and rheMacS SVs.

SV distribution were counts based on sliding windows (window size = 500 Kbp). Before plotting the distribution of bin counts with telomere distance, we identifies the centromeres of macaque in rheMac8 based on the results of (Rogers, et al. 2006), and the coordinates of each chromosomes’ centromeres were lifted over from rheMac2 to rheMac8.

SVs genotyping was conducted with Illumina deep sequencing data (HiSeq X Ten; PE read, >50-fold depth) in five unrelated Chinese rhesus monkeys using SVTyper (Chiang, et al. 2015). As insertion is unsupported for SVTyper, we obtained insertion genotypes by mapping WGS reads to rheMacS, which containing the novel insertion sequences in assembly. According to genotyped as deletion in the rheMacS-based alignments, we inferred insertion genotypes of each individual.

### Identification of ASSVs

Genome comparisons were performed using smartie-sv (Kronenberg, et al. 2018). We mapped each long-read assembled ape genome to the rheMacS assembly, including human-ZF1, chimpanzee-Clint_PTRv1 (CCP), gorilla-Susie3 (GS3) and orangutan-Susie_PABv1 (OSP) (Table S19). Each chromosome of rheMacS was mapped to each ape genome sequence for SV calling, which was referred to as forward calling. For the reverse calling, we called the SVs by mapping each chromosome of each ape back to the rheMacS genome (Figure S23). Then, we filtered the SVs by intersecting two SV sets and obtained the high-confident SVs for each genome pair. ASSVs were identified by overlapping all rheMacS–ape SV sets (Figure S11).

Given the high-quality annotation of the reference human genome, we also mapped rheMacS to GRCh38 (V38) to call SVs. The called ASSV sets based on ZF1 and GRCh38 were overlapped to obtain the SVs with GRCh38 coordinates for better annotation (Figure S11). Furthermore, to exclude the monkey-lineage-specific SVs, we used the published marmoset genome (assembly name: ASM275486; assembly ID: GCA_002754865.1) (Table S19).

### Annotation analysis for ASSVs

ASSV annotation was performed using VEP with the GRCh38 coordinates, and the upstream and downstream distances were both 5 Kbp.

### Simulation analysis of ASSVs

We generated a null distribution by randomly selecting a set of SVs as “ASSVs” from all rheMacS-GRCh38 SVs over many iterations (1 million times). We then assessed those SVs against the genes they intersect to determine how often genes are associated with one of the four traits (tail development, microcephaly, body size and adducted thumbs).

### Identification of ASSVs located in ADEs

The published ChIP-seq data was used to identify ADEs in eight brain regions (Xu, et al. 2018). We downloaded the 51,283 genomic regions with H3K27Ac modification signals, which were generated from three humans, two chimpanzees and three rhesus monkeys in each of the eight brain regions, including PFC, precentral gyrus (PcGm), occipital pole (OP), caudate nucleus (CN), putamen (Put), cerebellum (CB), white matter (WM), and thalamic nuclei (TN). We pair-wisely compared the counts of each enhancer among three species (macaque-human, macaque-chimpanzee and human-chimpanzee) and used the two-tailed unpaired t-test statistical assessment. We defined an enhancer as an ADE if this enhancer showed significant difference (P<0.05) of the same direction in both macaque-human and macaque-chimpanzee comparisons while showing no difference (P>0.05) in the human-chimpanzee comparison. According to this criterion, we obtained 7,155 ADEs (Table S25). To explore whether ASSVs occurred within these ADEs, we intersected the 7,155 ADEs and our ASSV set, and we found 87 ADEs disrupted by 111 ASSVs near 65 genes. To quantify these ADEs, we calculated the log_2_(fold-change) between apes (average of human and chimpanzee) and rhesus macaque. Functional annotation clustering of ASSVs was performed using DAVID (v6.8) (Huang, et al. 2009).

### Comparative gene expression analysis of neocortical layers

The gene expression data of neocortical layers in human, chimpanzee and macaque were obtained from the published data (He, et al. 2017), which contains samples from four humans, four chimpanzees and four macaques of neocortical layers, including 16 laminar sections (section 1S-16S) and part of the adjacent white matter (section 17S). The two-tail paired t-test was used in accessing expression differences between apes and macaques.

### Identification of ASSVs related to ape-specific traits

We collected the tail-development related-genes set from three sources: 1) genes related to caudal development and defects from the literature; 2) genes that showed tail-length changes (absent tail, short tail and long tail) and caudal vertebral body fusion phenotype in the mouse gene knockout models from the MGI database (http://www.informatics.jax.org/); and 3) genes with reportedly caudal regression syndrome or sacral agenesis in patients. An integrated table is shown in Table S27. The genes causing primary microcephaly were obtained from the previous study (Jayaraman, et al. 2018). The genes related to adducted thumbs were collected from Human Phenotype Ontology (https://hpo.jax.org/app/ ID: HP:0001181). The body size related genes were collected from the MGI database and literature that showed body size related changes (such as short stature and growth failure).

### ASSVs manual check and PCR validation

The sequences of the ASSV regions were extracted by SAMtools *view*, and sequence comparison was performed and plotted by MUMmer and LASTZ. The candidate ASSVs with 1 Kbp upstream/downstream sequences were aligned to the marmoset and gibbon genomes using MUMmer and blastn, and the best alignment score region was defined as the potential ASSV region in marmoset and gibbon. The ASSVs were classified into three categories based on manual check: 1) If the potential ASSV region and its 1 Kbp flanking sequences of marmoset could be completely aligned to the corresponding ASSV region of macaque using MUMmer and MEGA-X (Kumar, et al. 2018) (MUSCLE under default parameters), indicating that the regional sequences (SV + 1 Kbp flanking) were orthologous between marmoset and macaque. These ASSVs were defined as “high-confident” ASSVs; 2) If the potential ASSV region with 1 Kbp flanking sequences of marmoset could only be partially aligned to the corresponding ASSV sequence of macaque, we classified these ASSVs as “uncertain” ASSVs; and 3) If the potential ASSV region with 1 Kbp flanking sequence of marmoset could be completely aligned to the corresponding ASSV sequence of macaque except for the SV regions, suggesting that these SVs also exist in marmoset, we then defined these ASSVs as “false” ASSVs. The same criterion was employed when using the gibbon genome to distinguish ASSVs from GASSVs. To further evaluate the identified 17,000 ASSVs, we mapped all the ASSVs with 1 Kbp upstream/downstream sequences of rheMacS to the NGS marmoset genome using MUMmer and blastn with default parameters, and the high alignment score region (with a stringent cut off: blastn: E-value=0 and bit-score>1000; MUMmer: Sequence-identity>85%) was defined as the potential ASSV regions in marmoset. ASSVs below the threshold were considered as uncertain ASSVs. For PCR and Sanger sequencing validation, we included one rhesus macaque, one western hoolock gibbon (*Hoolock hoolock*), one chimpanzee and one human. The primers were designed by Primer Premier5, and the extended 100 bp sequences were included at the ASSV breakpoints as the PCR targets. The PCR products were visualized by agarose gel electrophoresis to verify the lengths of ASSVs and for Sanger sequencing. All the presented SVs in this study were validated both by PCR and Sanger sequencing.

### Ethics Statement

The monkey was housed at the AAALAC (Association for Assessment and Accreditation of Laboratory Animal Care) accredited facility of Primate Research Center of Kunming Institute of Zoology. The animal protocol was approved in advance by the Institutional Animal Care and Use Committee of Kunming Institute of Zoology (Approval No: SYDW-2010002).

### Data Availability

The PacBio sequence data, Illumina sequencing reads, rheMacS assembly and its annotation files have been deposited in NCBI and GSA (Genome Sequence Archive) under the project accession numbers of PRJNA514196 and PRJCA001197, respectively.

## Author contributions

B.S., X.Q., X.L. and Y.H. designed the project; X.L., T.H., X.M. collected the samples; Y.H., B.Z., J.J., X.M., Z.N.K., P.A., Y.G. and Y.Y. performed bioinformatics analyses; T.H., B.Z., and Y.H. performed genotyping and sequencing experiments; B.S. and Y.H. wrote the manuscript. All authors discussed the results and implications and commented on the manuscript.

## Acknowledgements

This study was supported by grants from the Strategic Priority Research Program of the Chinese Academy of Sciences (XDB13010000), and the National Natural Science Foundation of China (31730088 and 31621062). This work was supported, in part, by US National Institutes of Health (NIH) grant R01HG002385 to E.E.E. E.E.E. is an investigator of the Howard Hughes Medical Institute.

## Supplementary Figures

**Figure S1.**
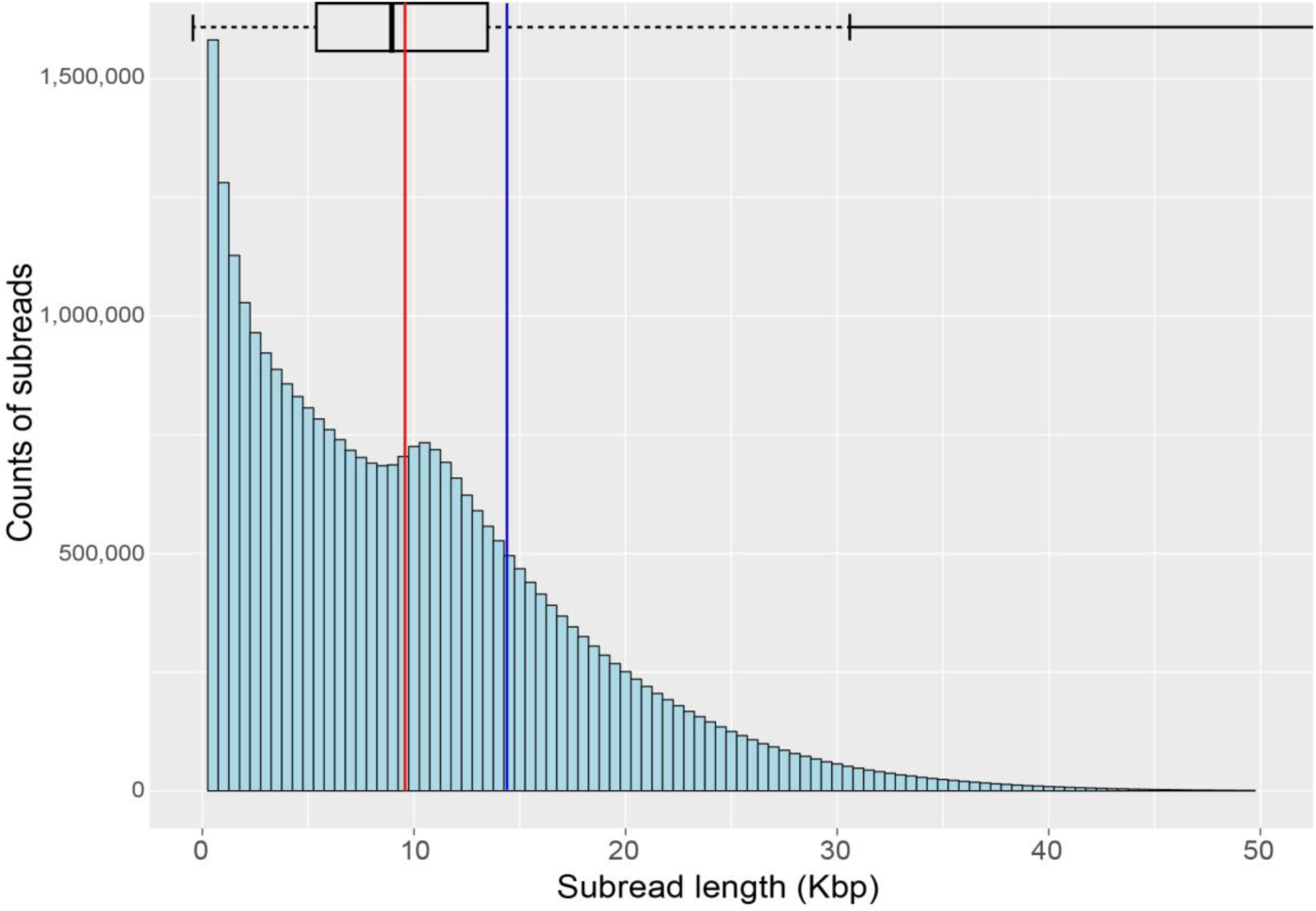
Distribution of subread length of rheMacS. Marginal box plot indicates quartiles. The mean subread length is 9.7 Kbp (red line) and the N50 subread length is 14.7 Kbp (blue line).

**Figure S2.**
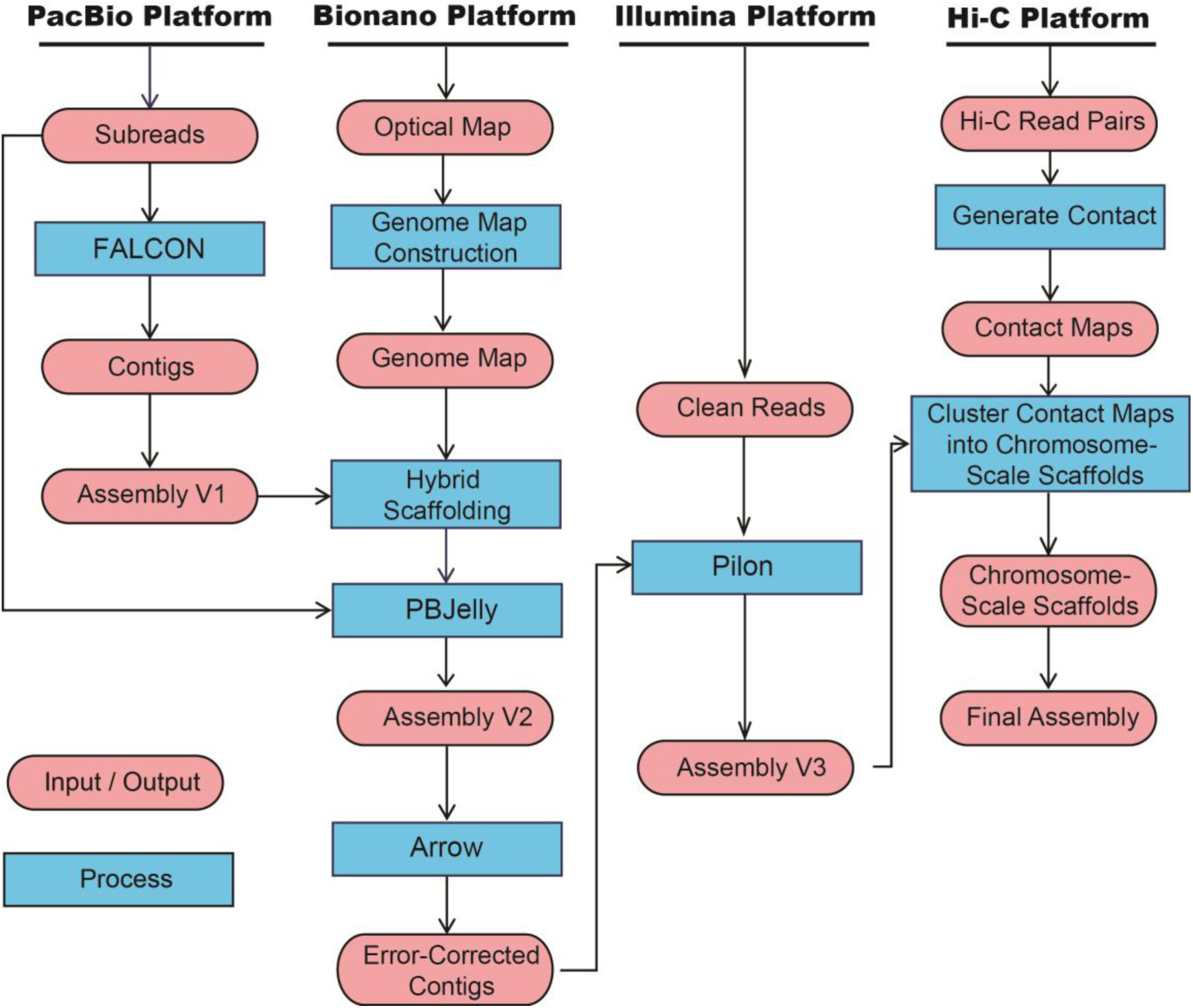
Data generation and *de novo* assembly pipeline.

**Figure S3.**
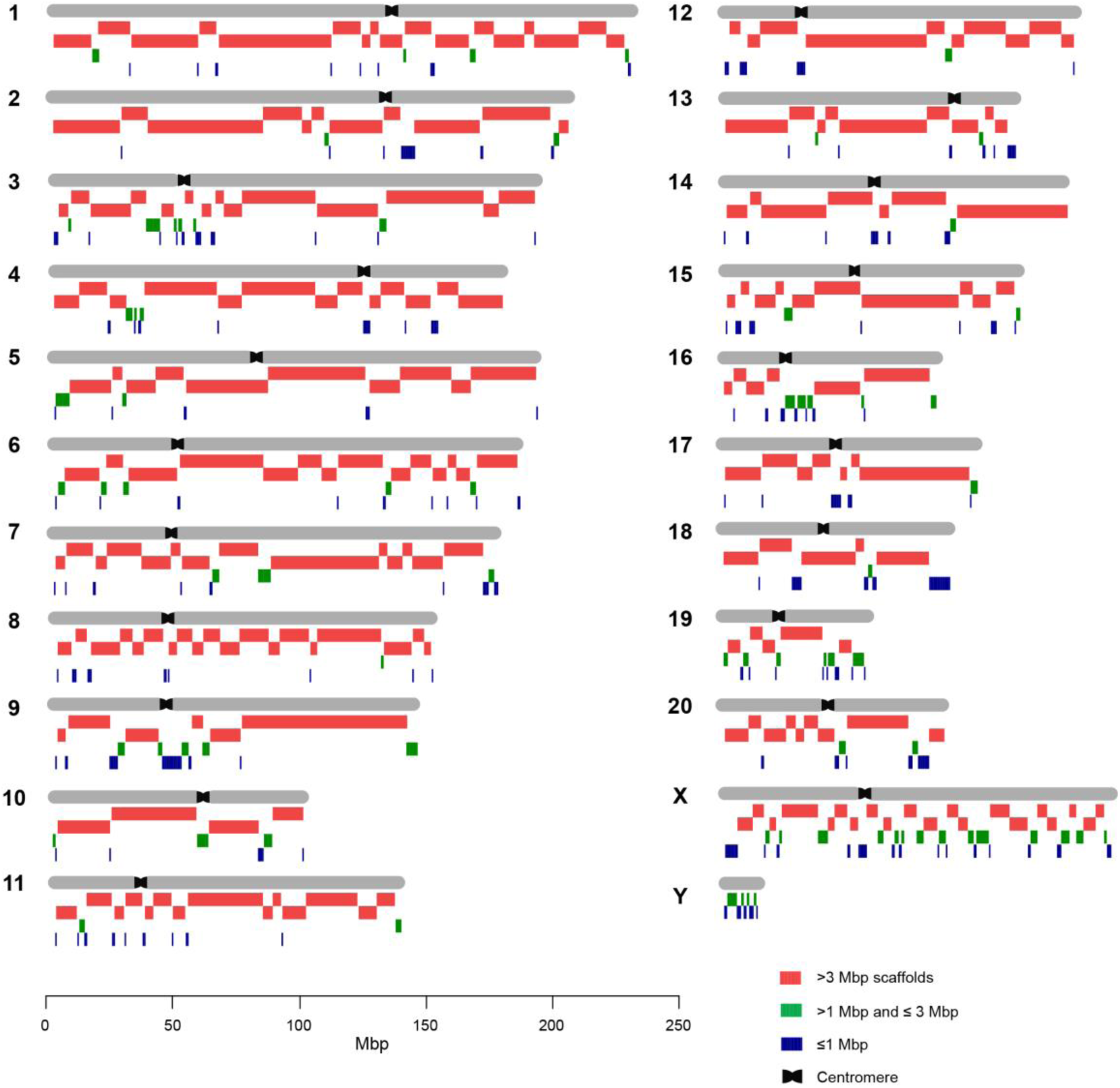
The chromosomal distribution of scaffolds of the rheMacS genome assembly. The red rectangles represent scaffolds >3 Mbp; green and blue rectangles correspond to the scaffolds with lengths between 1 Mbp and 3 Mbp and those <1 Mbp, respectively. The centromeres of each chromosomes are indicated according to (Rogers, et al. 2006).

**Figure S4.**
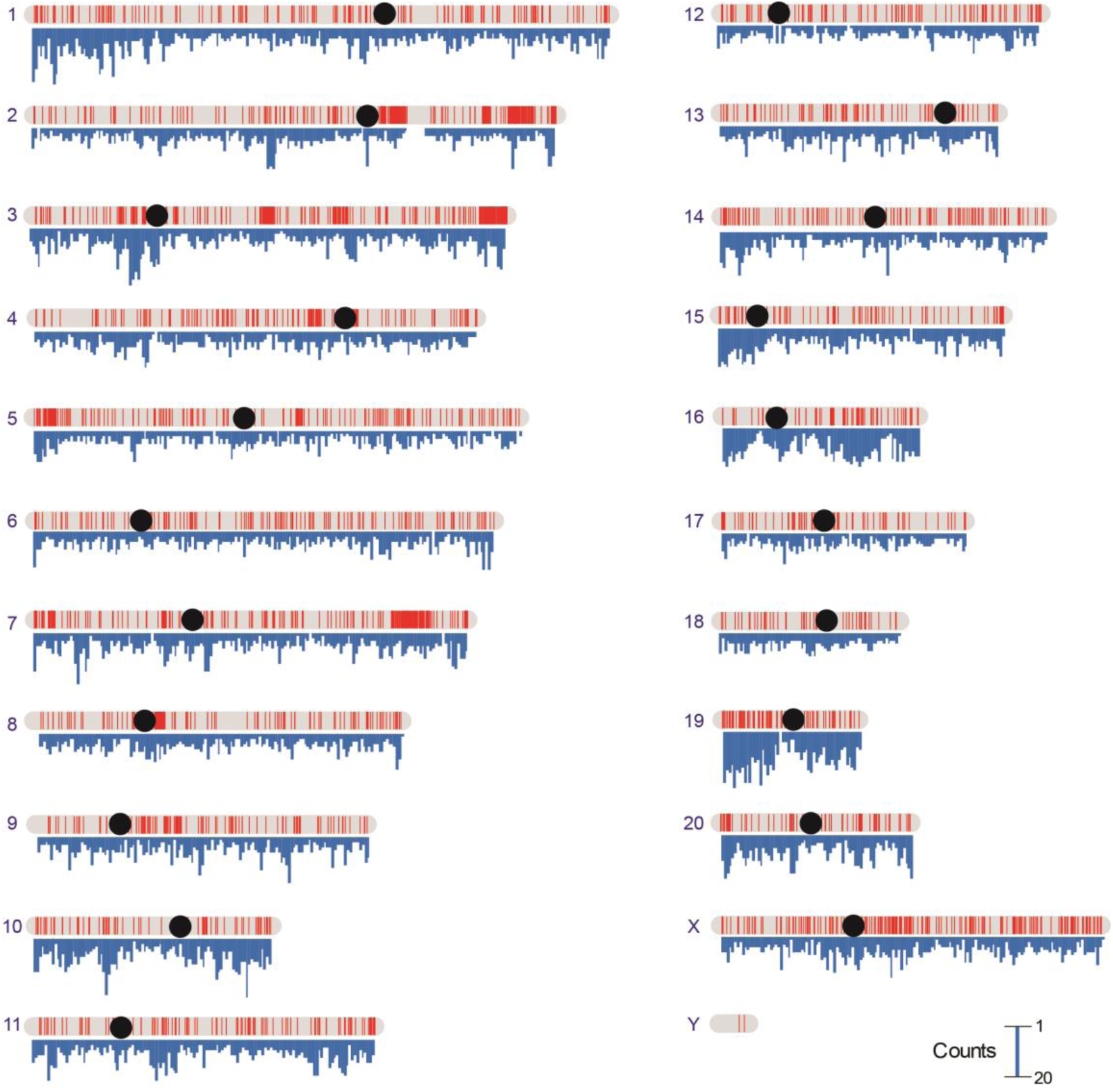
Distribution of closed gaps in rheMac8 (only the >1 Kbp closed gaps are shown). The blue-colored histograms on the axes of each chromosome indicate the counts of the closed gaps based on 500 Kbp windows. The black dot on each chromosome indicates the centromere position.

**Figure S5.**
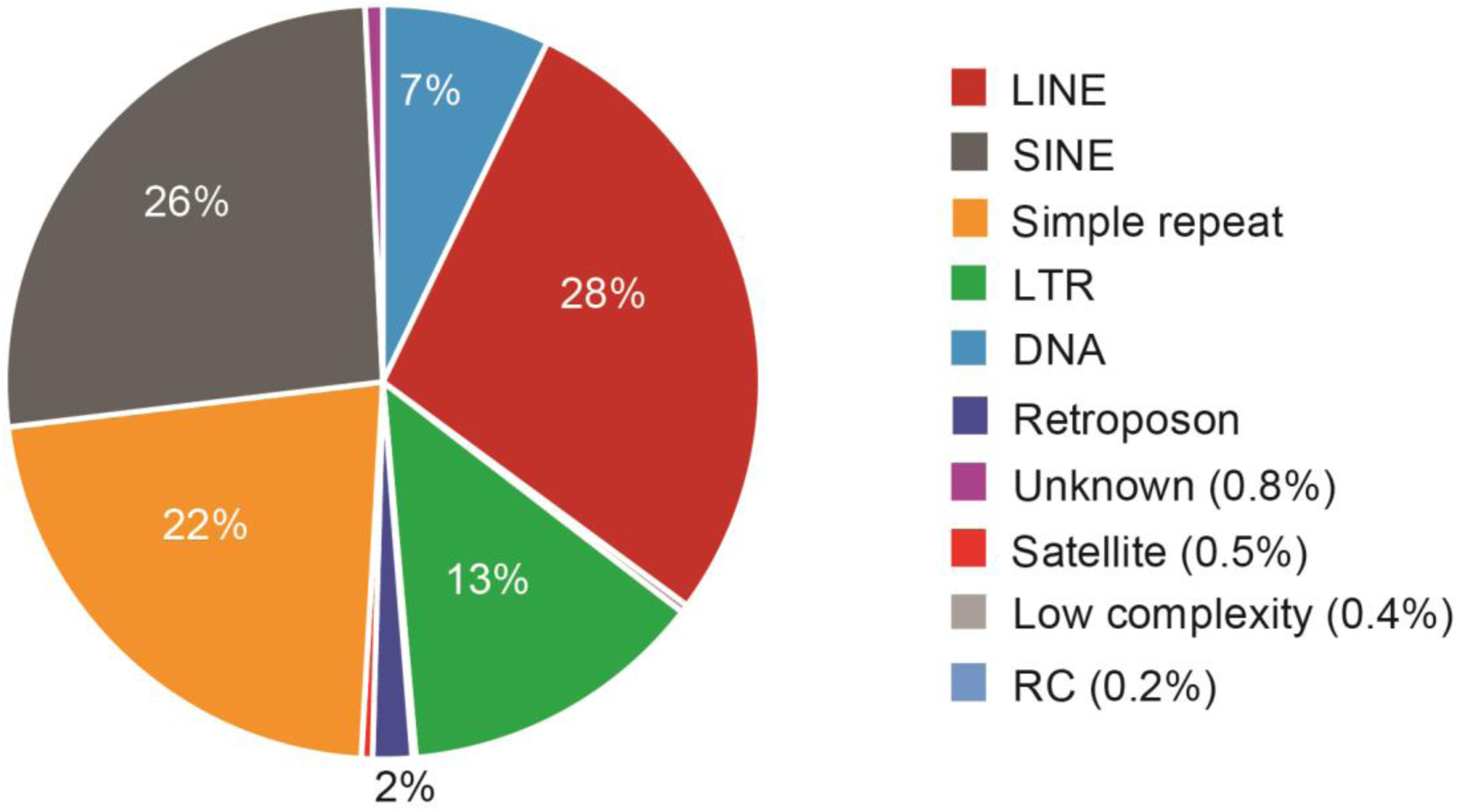
Repeat annotation for the closed gaps in rheMac8. LINE: long interspersed nuclear elements; SINE: short interspersed nuclear elements; LTR: long terminal repeats; RC: rolling-circle transposition.

**Figure S6.**
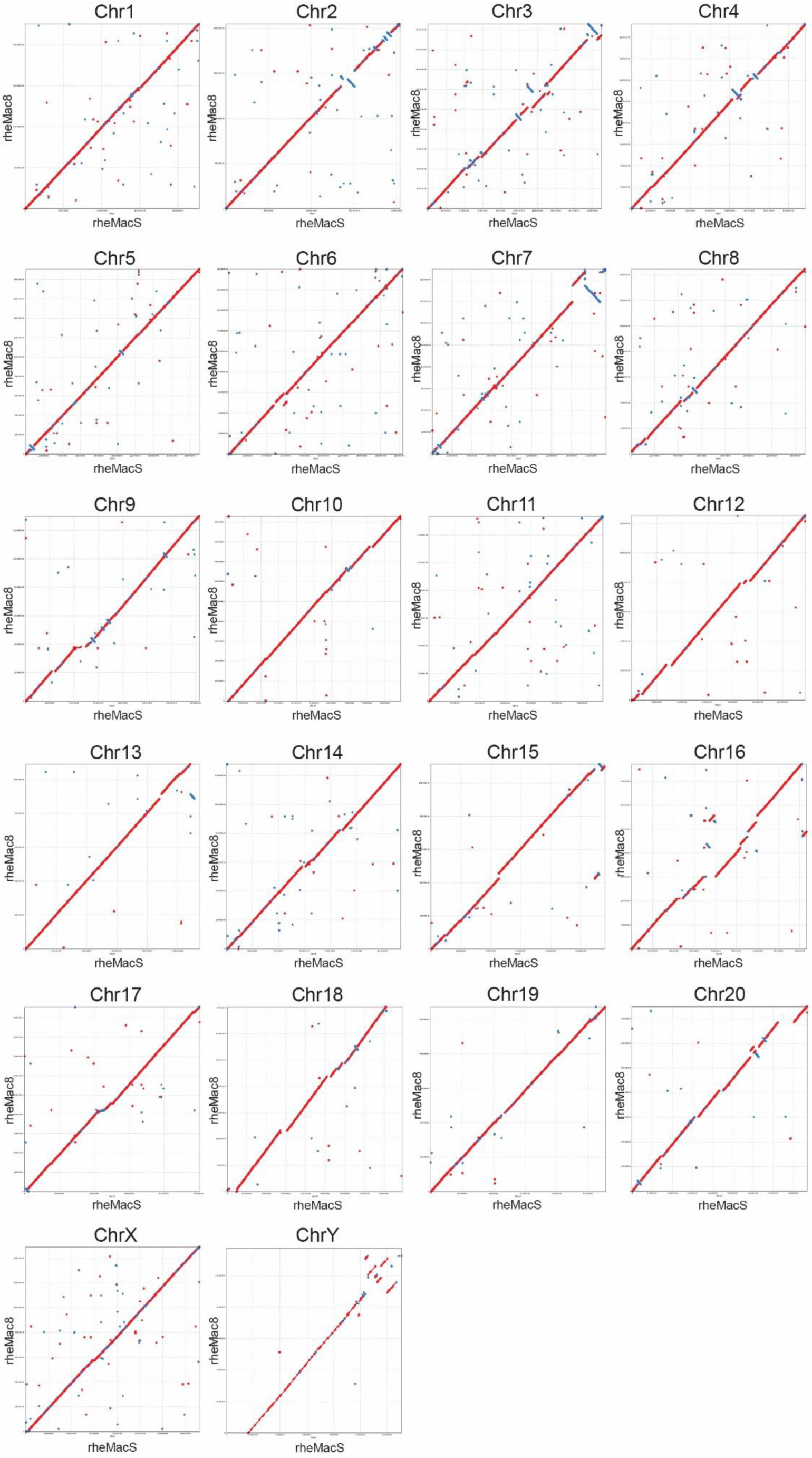
Dot plots of assembly comparison between rheMacS and rheMac8.

**Figure S7.**
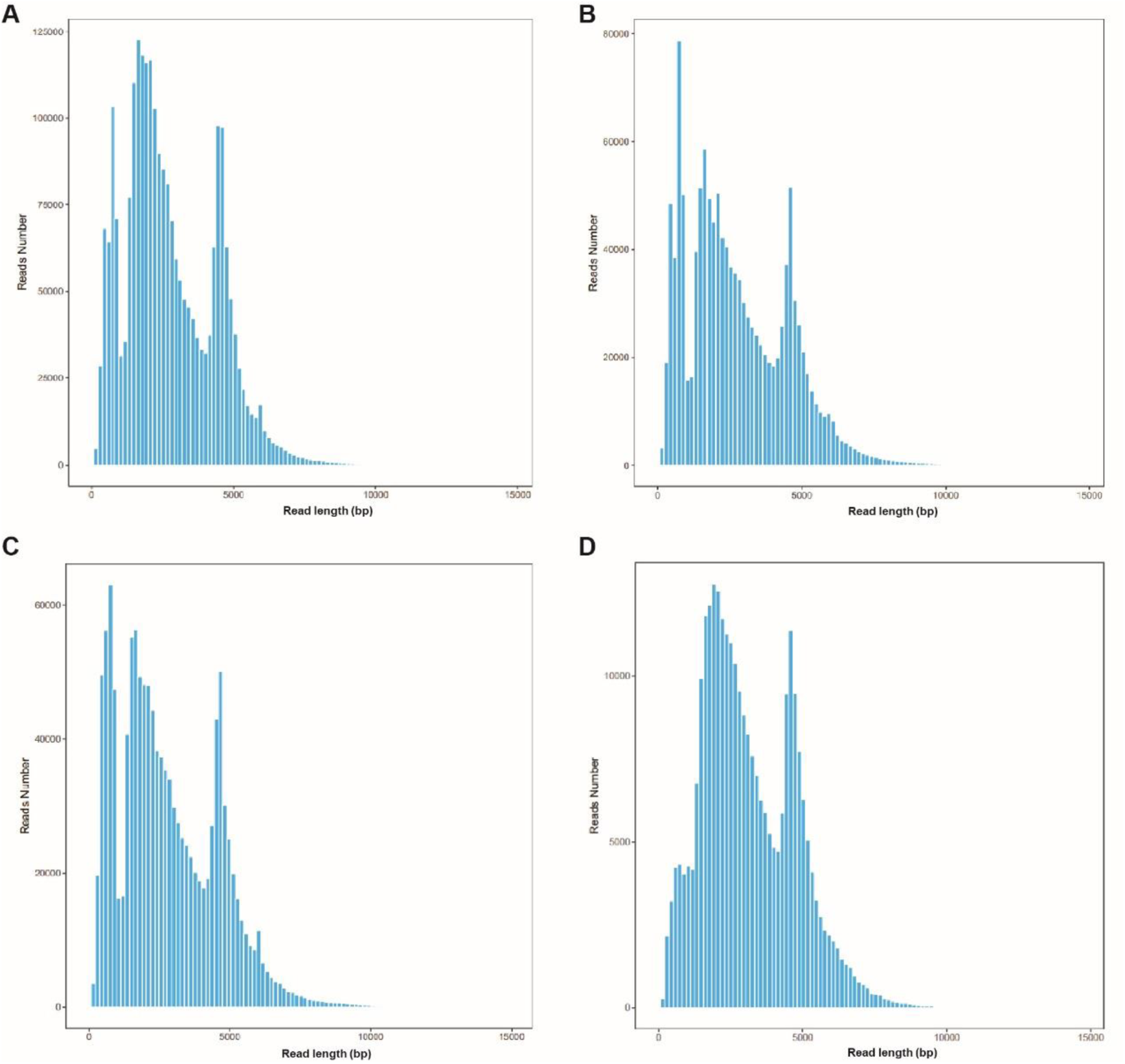
Distribution of transcripts. **(A)** Length distribution of Full-length non-chimeric (FLNC) reads. **(B)** Length distribution of consensus sequences. **(C)** Length distribution of the NGS-corrected reads. **(D)** Length distribution of collapsed isoforms.

**Figure S8.**
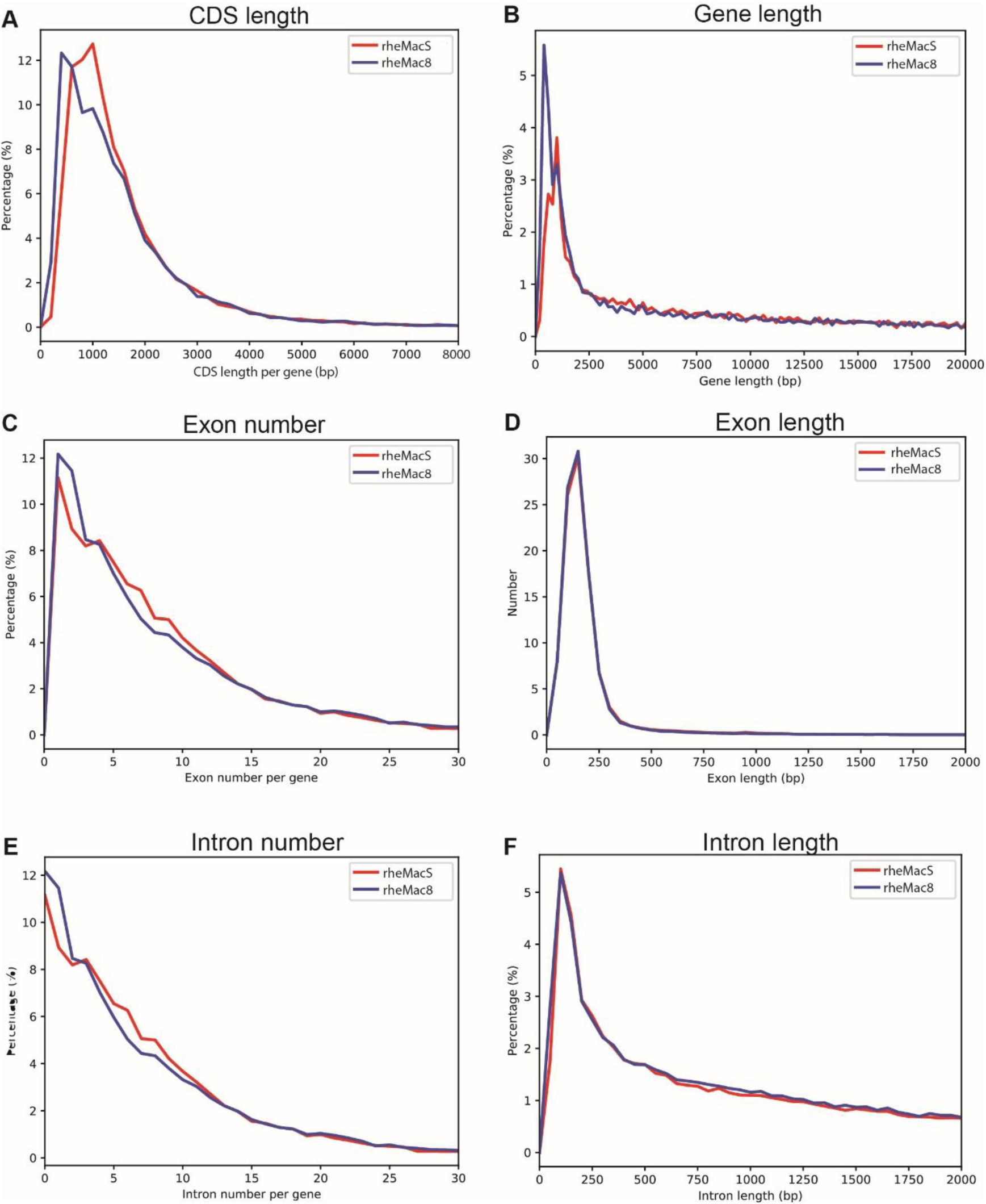
Comparison of gene structures between rheMacS and rheMac8.

**Figure S9.**
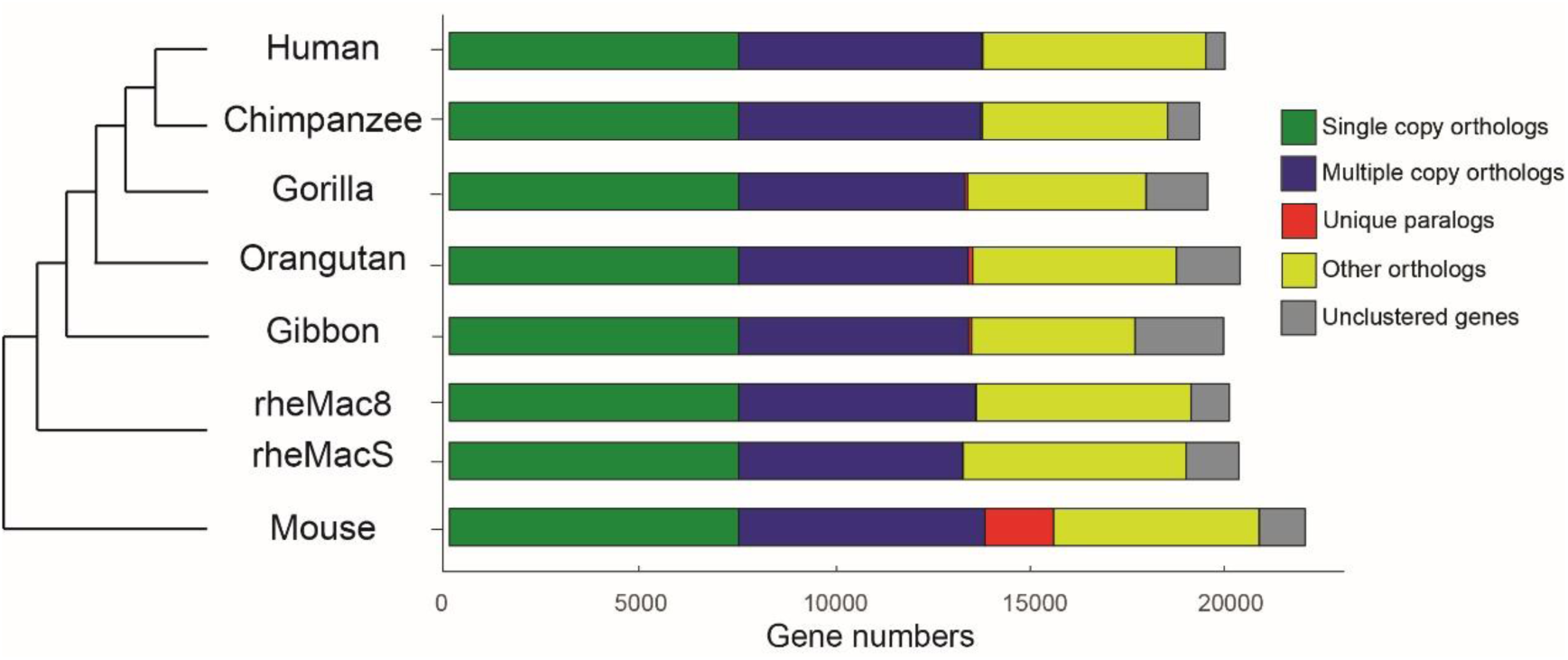
Comparison of orthologous gene families among rheMacS, rheMac8 and genome assemblies of other species.

**Figure S10.**
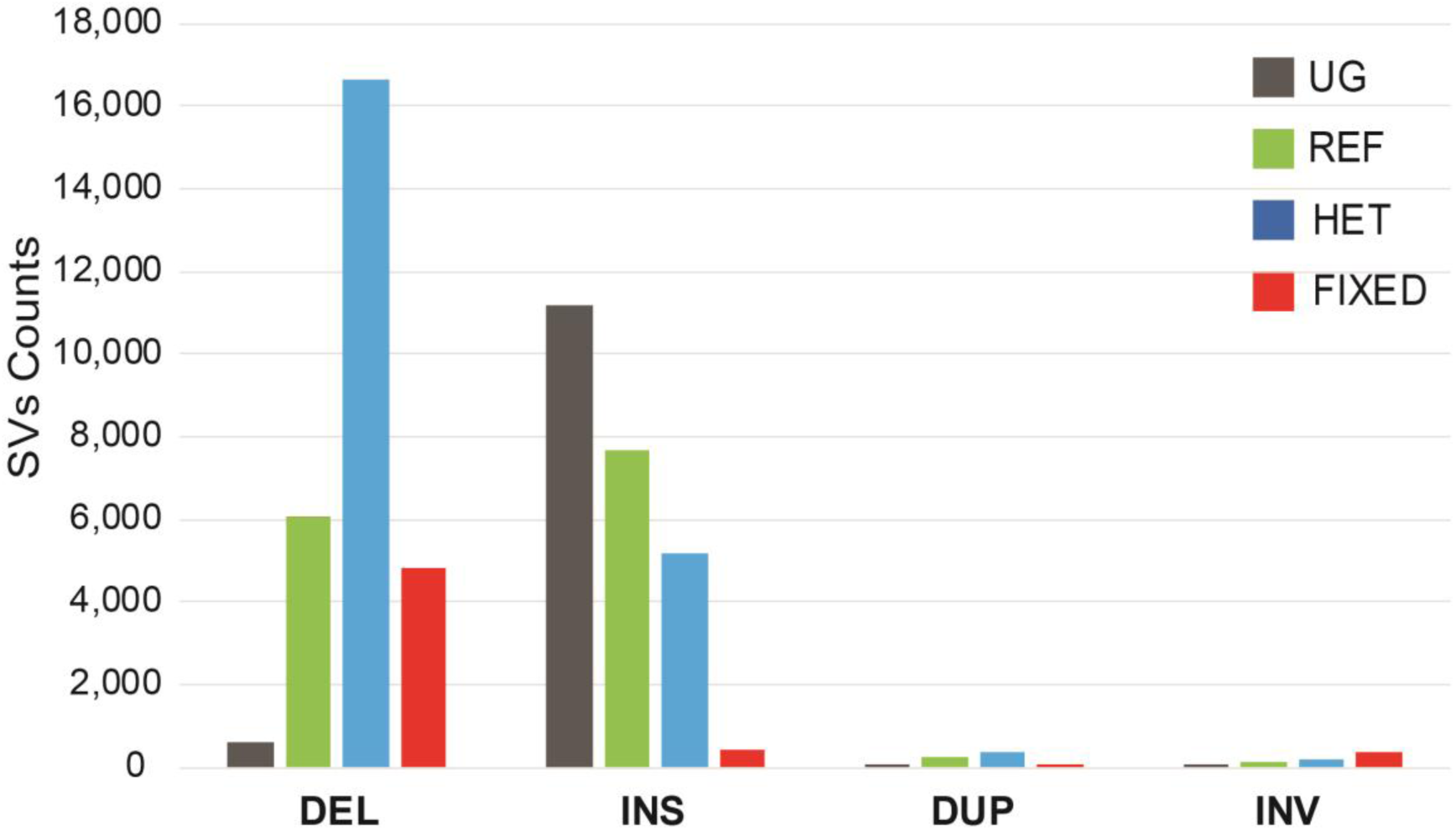
Genotyping results of 53,916 SVs of rheMacS in five unrelated Chinese rhesus monkeys. UG: ungenotyped SVs; REF: reference SVs; HET: heterozygous SVs; FIXED: fixed SVs.

**Figure S11.**
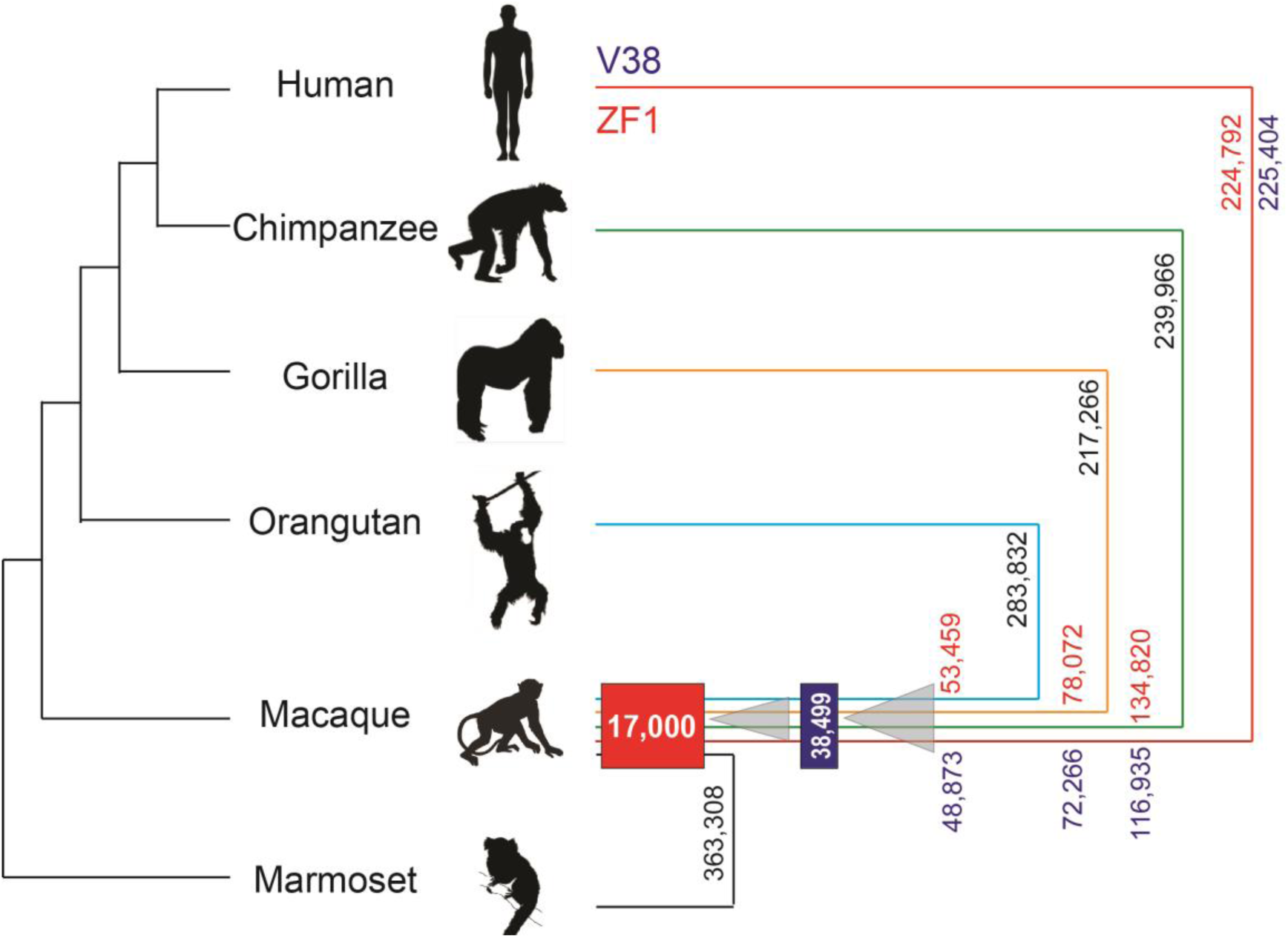
Schematic diagram of ASSV calling. The ape and monkey cladogram show SVs assigned to lineages according to the assembly comparison. Gibbon is excluded due to its poor assembly quality. The SV count is shown on each branch. For human genomes, a long-read assembled genome ZF1 and the reference genome GRCh38 are used for SV calling, and the SVs number are marked in red (ZF1) and blue (V38). The common marmoset genome assembled by short-read sequencing is used to exclude the SVs that occurred in the monkey lineage. After obtaining the 38,499 SVs between macaque and ape, we then characterized SVs between macaque and marmoset and obtained 363,308 SVs. An SV is defined as an ape-specific structural variant or ASSV if it is included in the 38,499 set, but not in the 363,308 set. We obtained 17,000 ASSVs in total. The final set of ASSVs are highlighted in red box.

**Figure S12.**
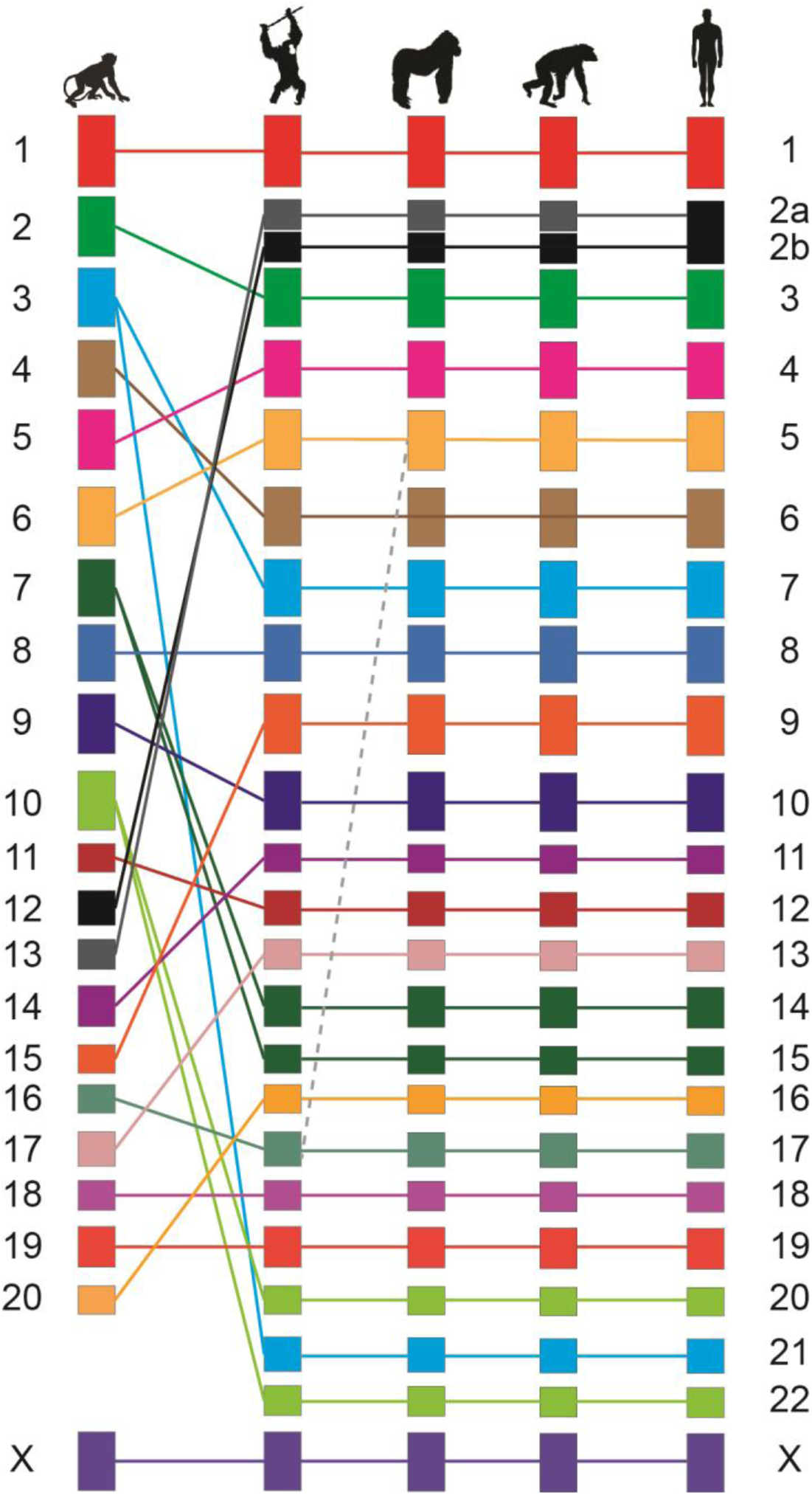
The orthologous locations of ASSVs on each chromosome between rhesus macaque and apes are consistent with the known chromosomal synteny. The dashed line refers to the known translocation event between chromosome 17 and chromosome 5 in gorilla.

**Figure S13.**
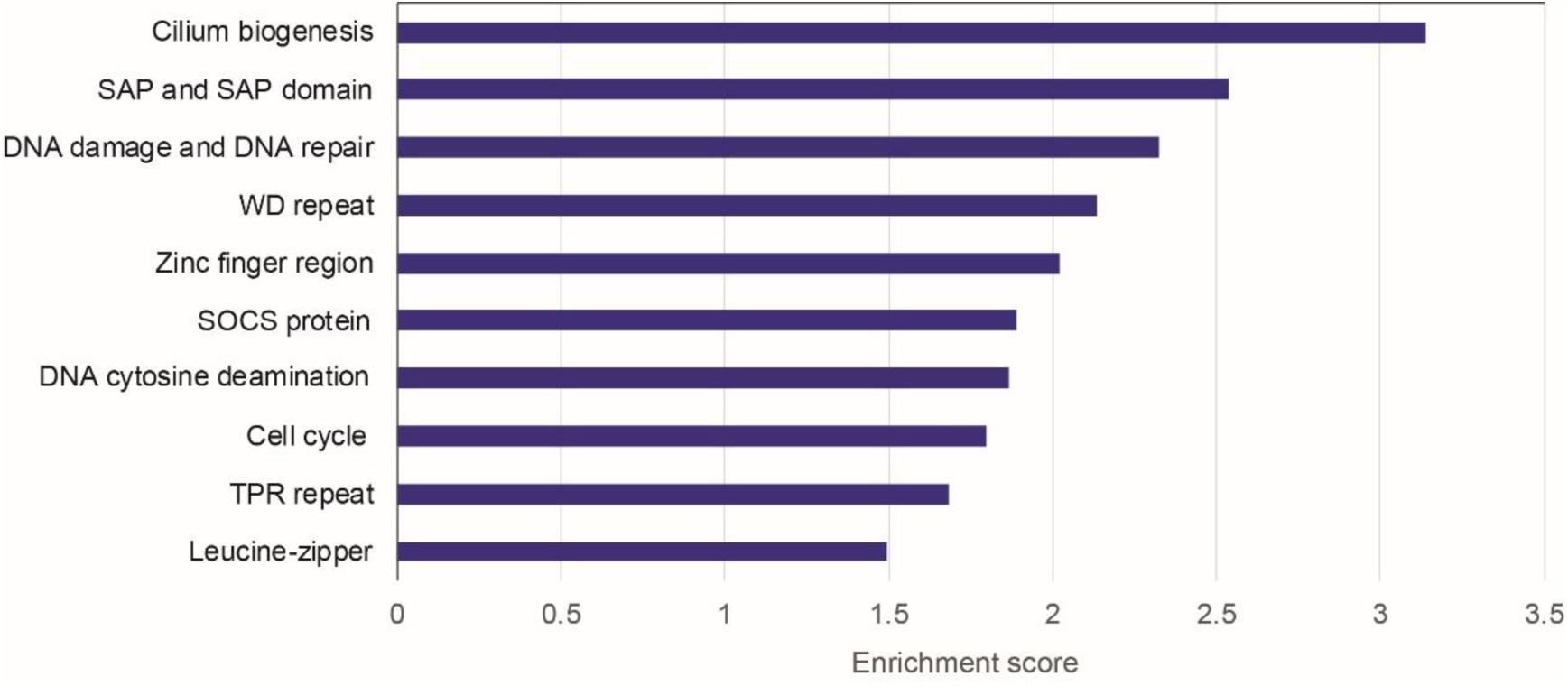
Functional enrichment of the 3,412 ASSV-related genes. The top 5% (10/208) categories are shown.

**Figure S14.**
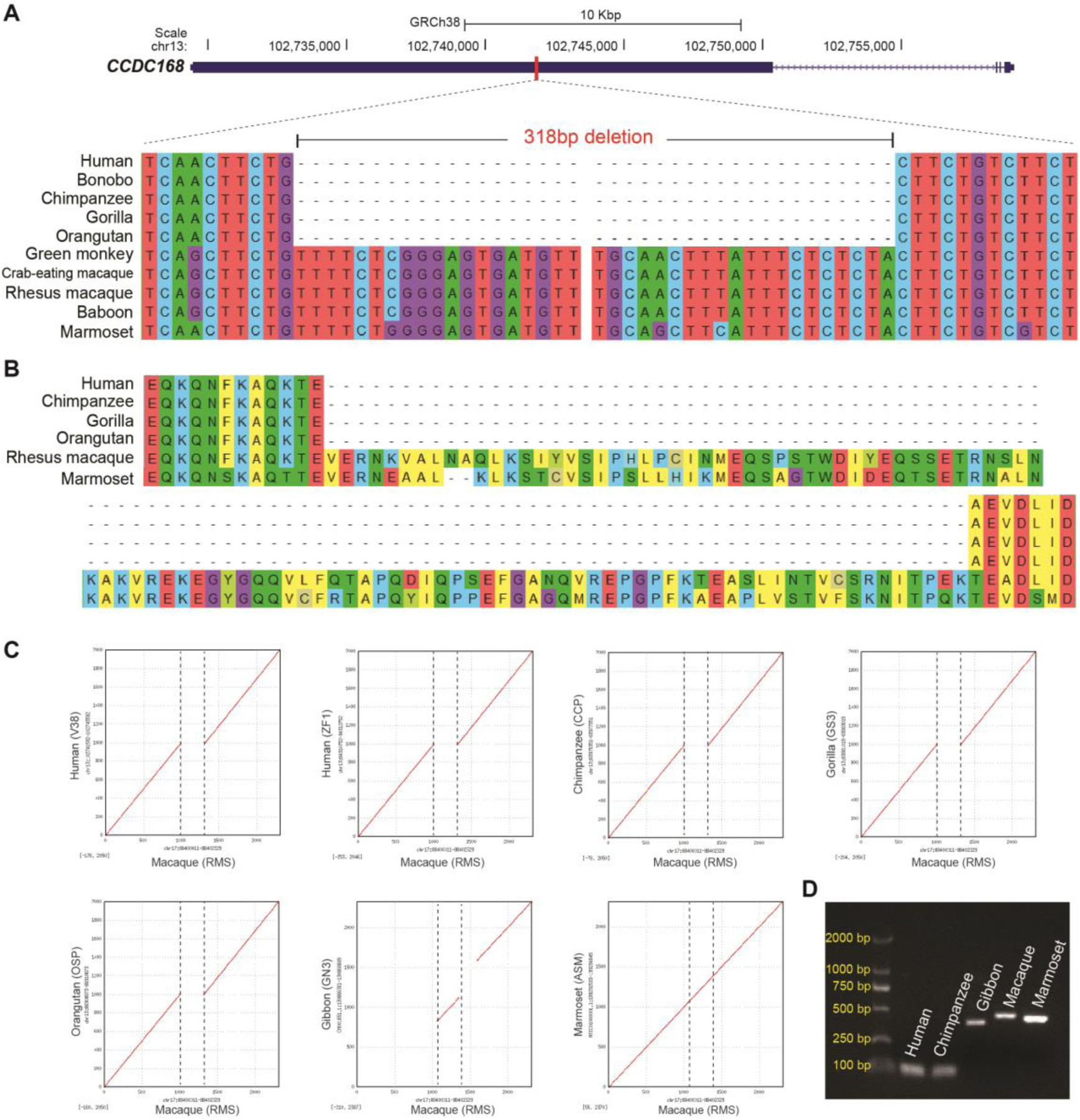
An ASSV located in gene-coding regions. **(A)** A 318 bp deletion located in *CCDC168* in the great ape lineage. The graph shows multiple comparative alignments of the 318 bp deletion region. **(B)** Amino-acid alignments. The 318 bp coding-region deletion leads to a 106-amino acid deletion in the ape lineage. **(C)** Dot plots for pairwise comparison of the 318 bp deletion region (1 Kbp downstream and upstream flanking sequences) between macaque and apes. **(D)** PCR validation for 318 bp deletion in *CCDC168*.

**Figure S15.**
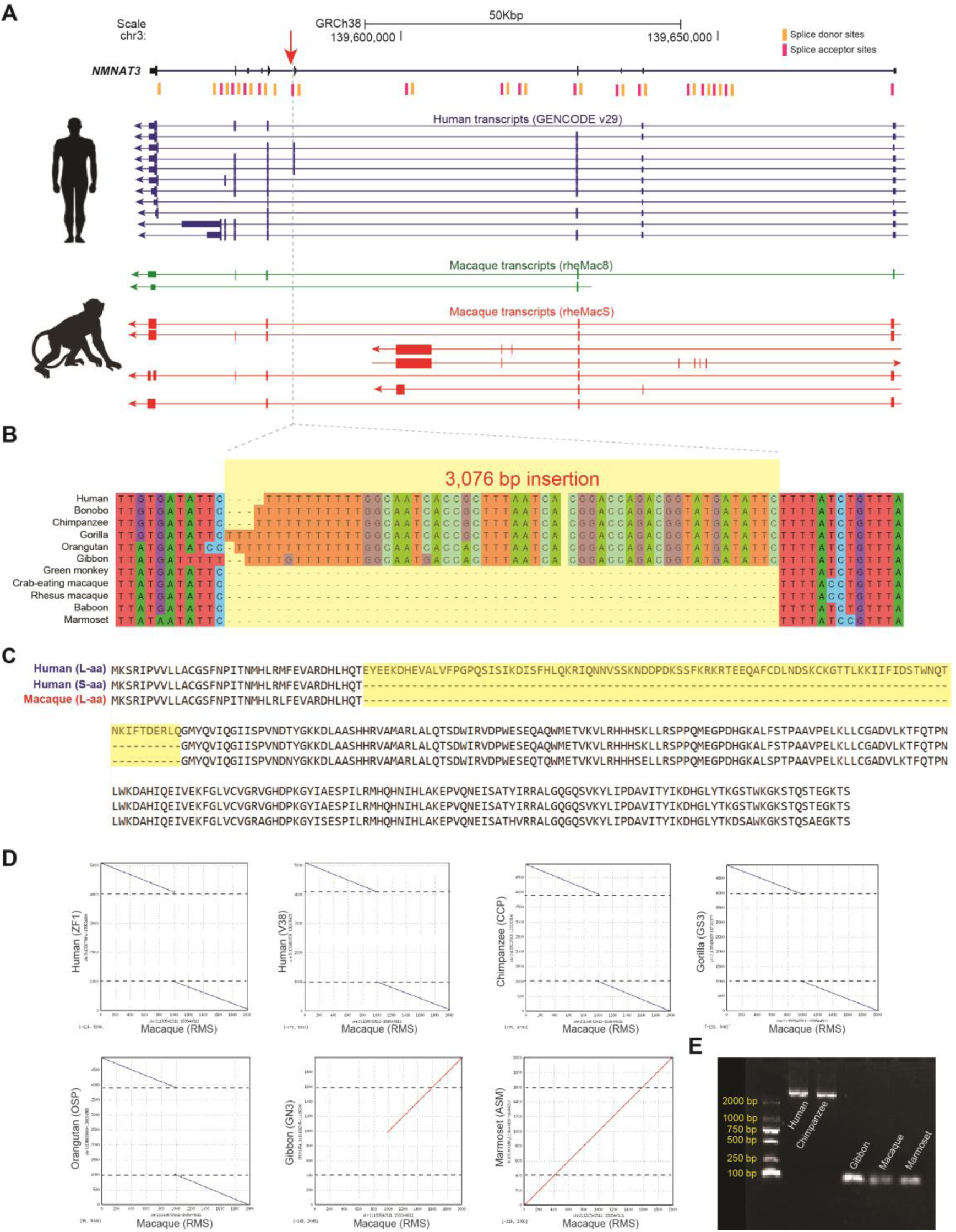
A 3,076 bp insertion located in the splice acceptor of *NMNAT3* in the great ape lineage. **(A)** The genomic location and transcript isoform comparison of *NMNAT3* between human and macaque. **(B)** Multiple comparative alignments of the 3,076 bp insertion region among primates. **(C)** Amino-acid alignment of *NMNAT3* between human and macaque. A human-specific protein isoform (highlighted) is identified, which is caused by the 3,076 bp insertion. **(D)** Dot plot for pairwise comparison of the insertion region (1 Kbp downstream and upstream flanking sequences) between macaque and apes. **(E)** PCR validation for 3,076 bp insertion in *NMNAT3*.

**Figure S16.**
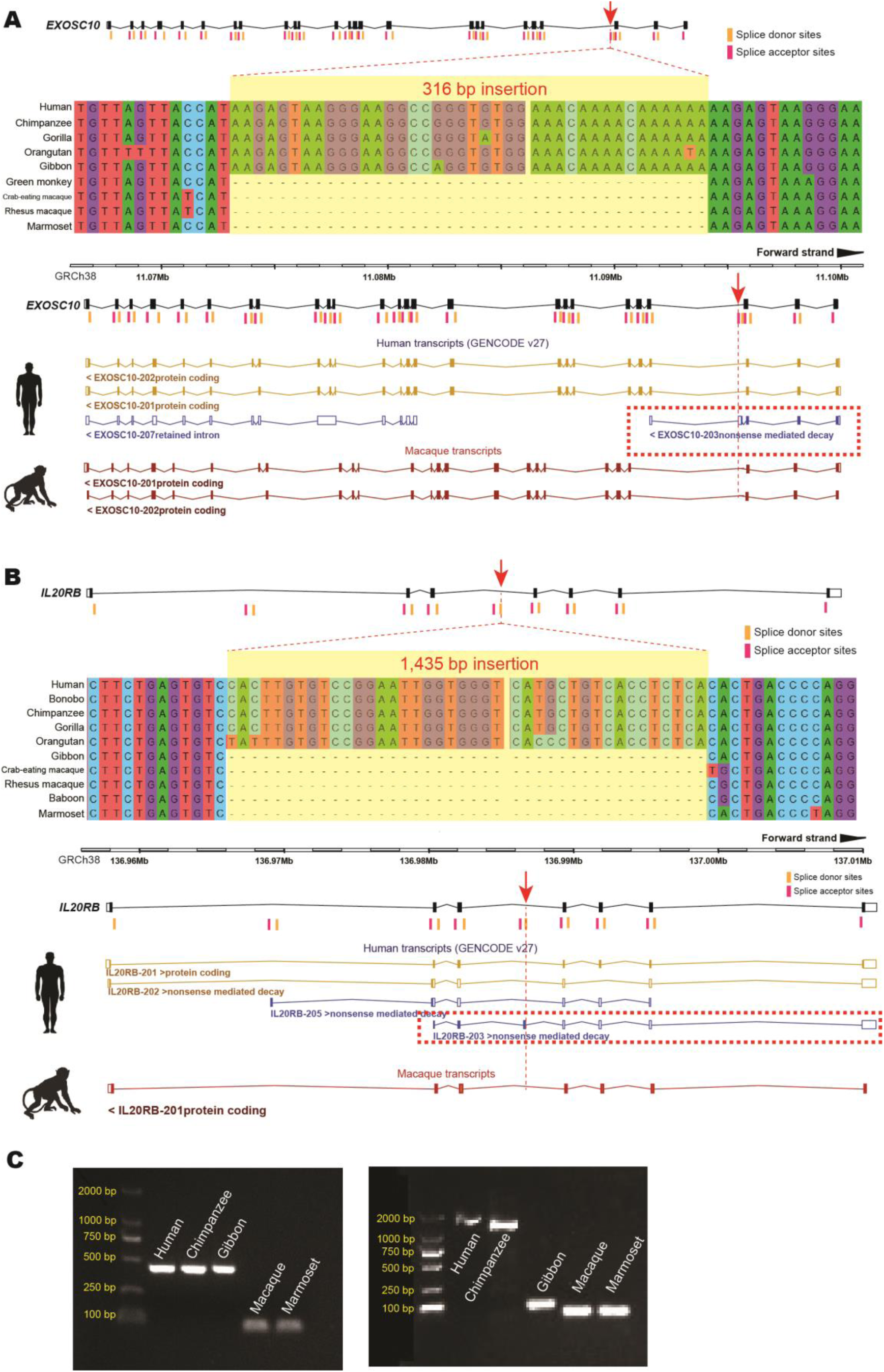
Two ASSVs located at the splice sites of *EXOSC10* and *IL20RB.* **(A)** A 316 bp insertion located in the splice acceptor of *EXOSC10*, resulting in a nonsense mediated decay (NMD) transcript (dashed frame) in the human lineage. The genomic location and multiple comparative alignments of the 316 bp insertion region (up-panel); transcript comparison of *EXOSC10* between human and macaque. **(B)** A 1,435 bp insertion located in the splice donor of *IL20RB*, resulting in an NMD transcript (dashed frame) in the human lineage. The genomic location and multiple comparative alignments of the 1,435 bp insertion region (up-panel); transcripts comparison of *IL20RB* between human and macaque. **(C)** PCR validations of 316 bp insertion in *EXOSC10* (left) and 1,435 bp insertion in *IL20RB* (right).

**Figure S17.**
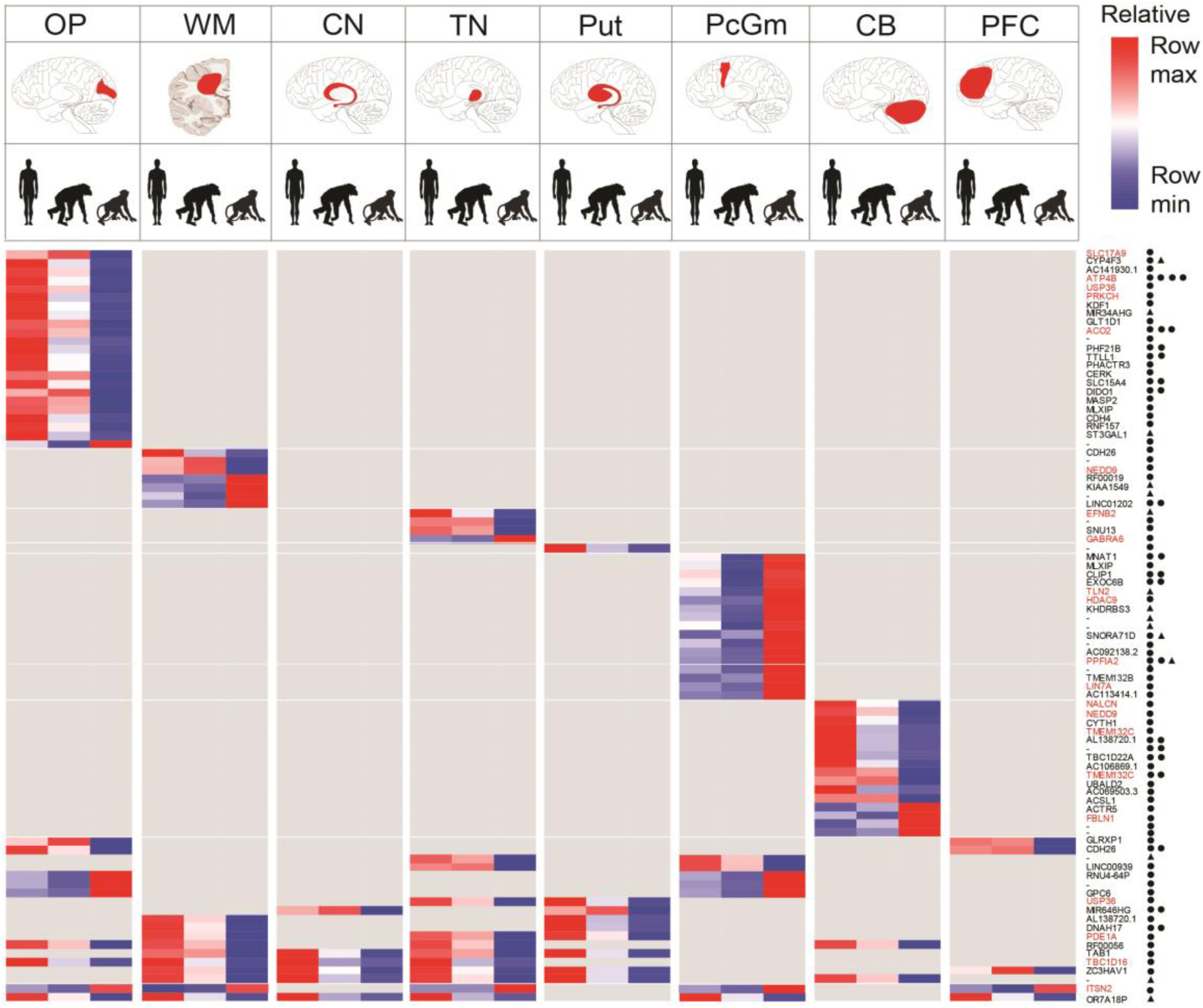
Heatmap for ADEs with candidate ASSVs in eight brain regions. There are 87 ADEs (111 candidate ASSVs). The nearest genes are indicated and the corresponding brain regions are shown in red. The neuro-function-related genes are highlighted (red). ASSV deletions (circles) and insertions (triangles) are denoted.

**Figure S18.**
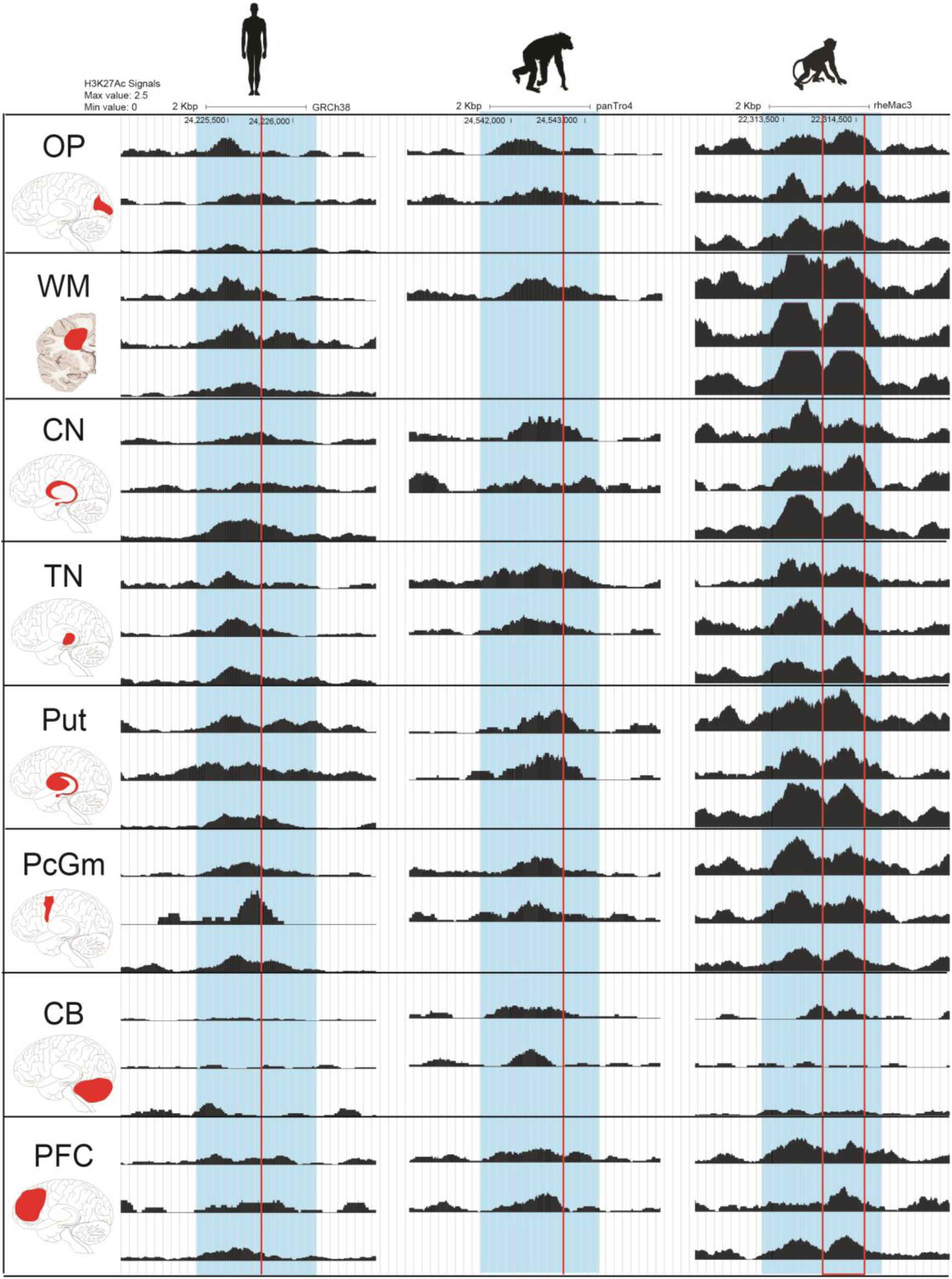
Comparison of H3K27Ac signals of the ADE between apes and macaques, which possess an ASSV (587 bp deletion) in *ITSN2*. It shows that the ADE exhibits significant difference between apes (human: n=3 and chimp: n=2) and macaques (n=3) in five brain regions. Shadow in light blue refers to ADE region, vertical lines and box in red refer to the ASSV region.

**Figure S19.**
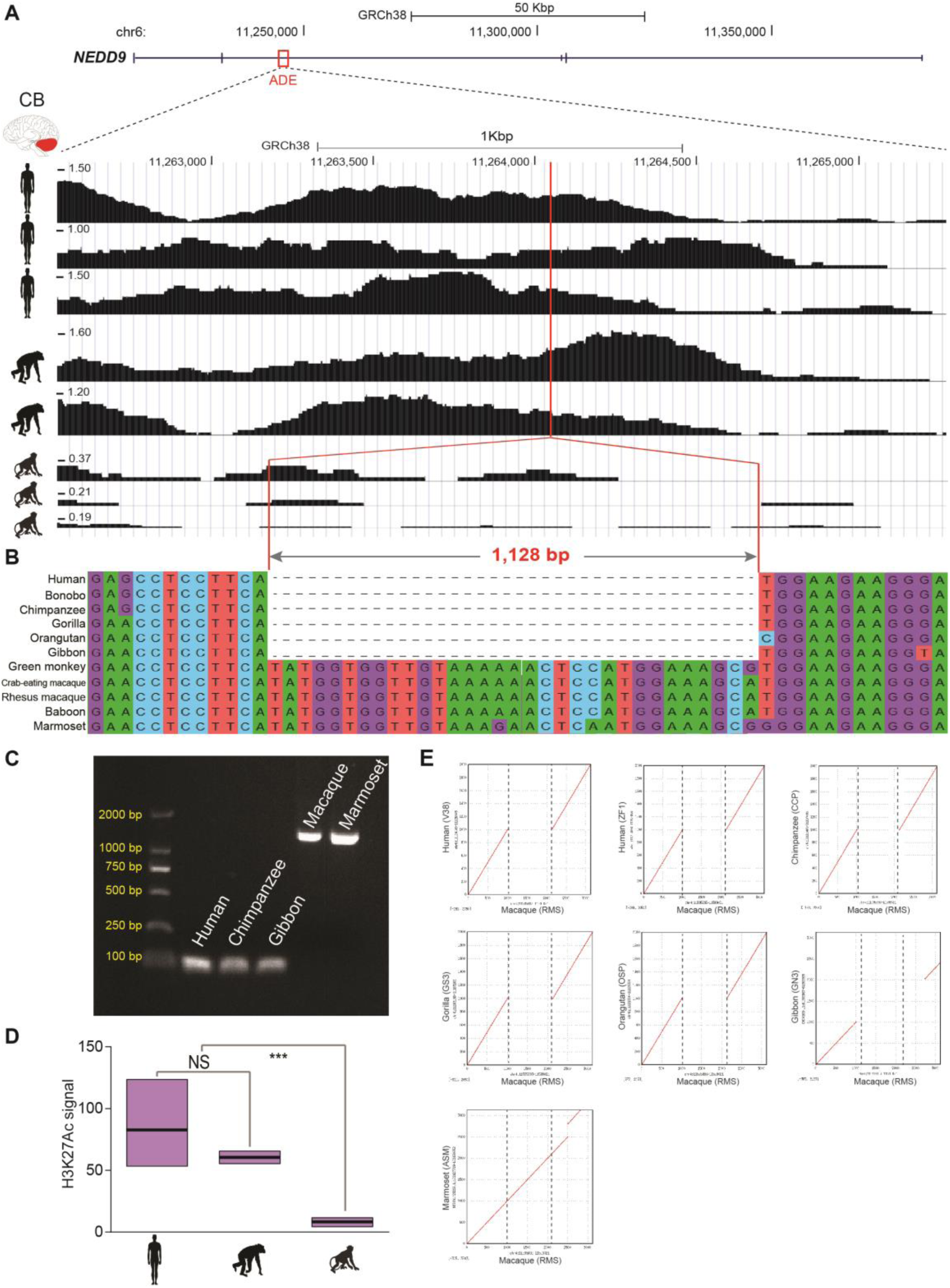
A 1,128 bp deletion located at an ADE of *NEDD9* in the ape lineage. **(A)** Location of the 1,128 bp deletion in *NEDD9* and the H3K27Ac signals among human, chimpanzee and macaque. **(B)** Sequence alignment of the deletion region among apes and macaque. **(C)** PCR validation. **(D)** Comparison of the H3K27Ac signals among human, chimpanzee and macaque. (***-P<0.001; NS-not significant, P>0.05) **(E)** Dot plots for pairwise comparison of the 1,128 bp deletion region (1 Kbp downstream and upstream flanking sequences) between macaque and apes.

**Figure S20.**
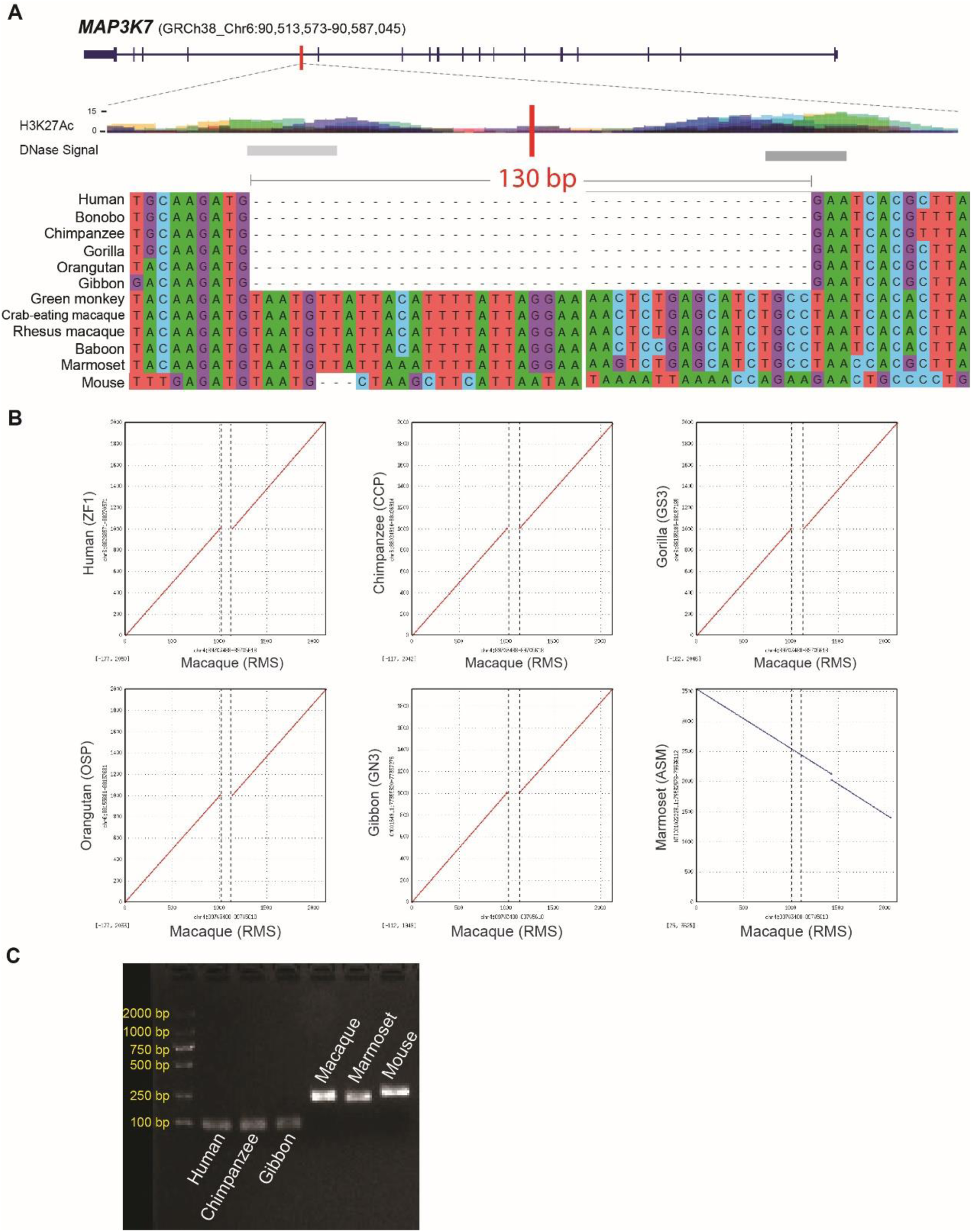
An ASSV (130 bp deletion) located in a tail-development-related gene *MAP3K7*. **(A)** Location of the 130 bp deletion of *MAP3K7* in apes and sequence alignments. **(B)** Dot plot for the pairwise comparison of the 130 bp deletion region (1 Kbp downstream and upstream flanking sequences) between macaque and apes. **(C)** PCR validation of the 130 bp deletion.

**Figure S21.**
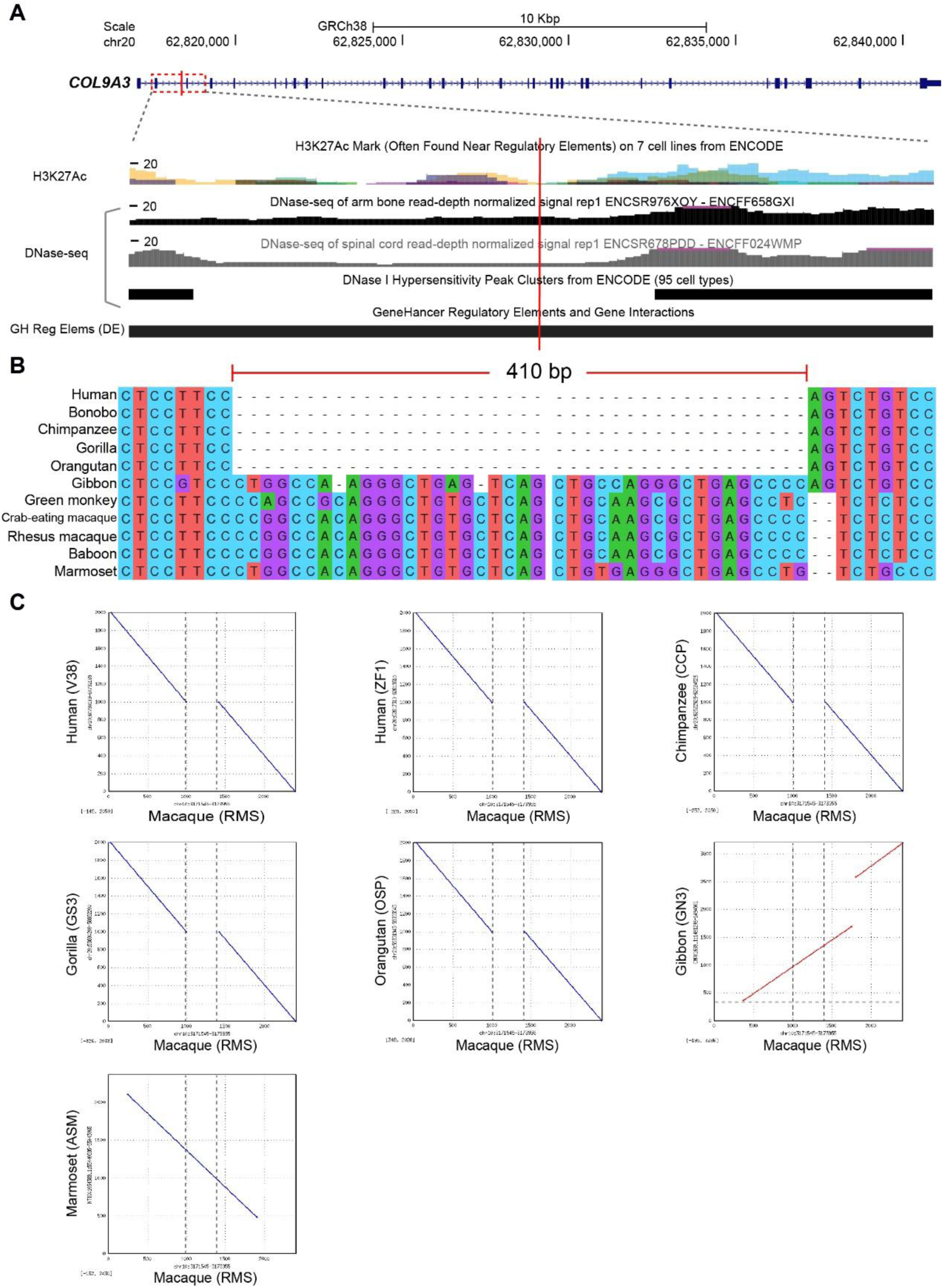
A GASSV (410 bp deletion) located in the intron region of *COL9A3*, a gene related to body size. **(A)** Location of the 410 bp deletion in great apes and the gene regulatory annotations (from ENCODE). **(B)** Sequence alignments among primates. **(C)** Dot plot for the pairwise comparison of the deletion region (1 Kbp downstream and upstream flanking sequences) between macaque and apes.

**Figure S22.**
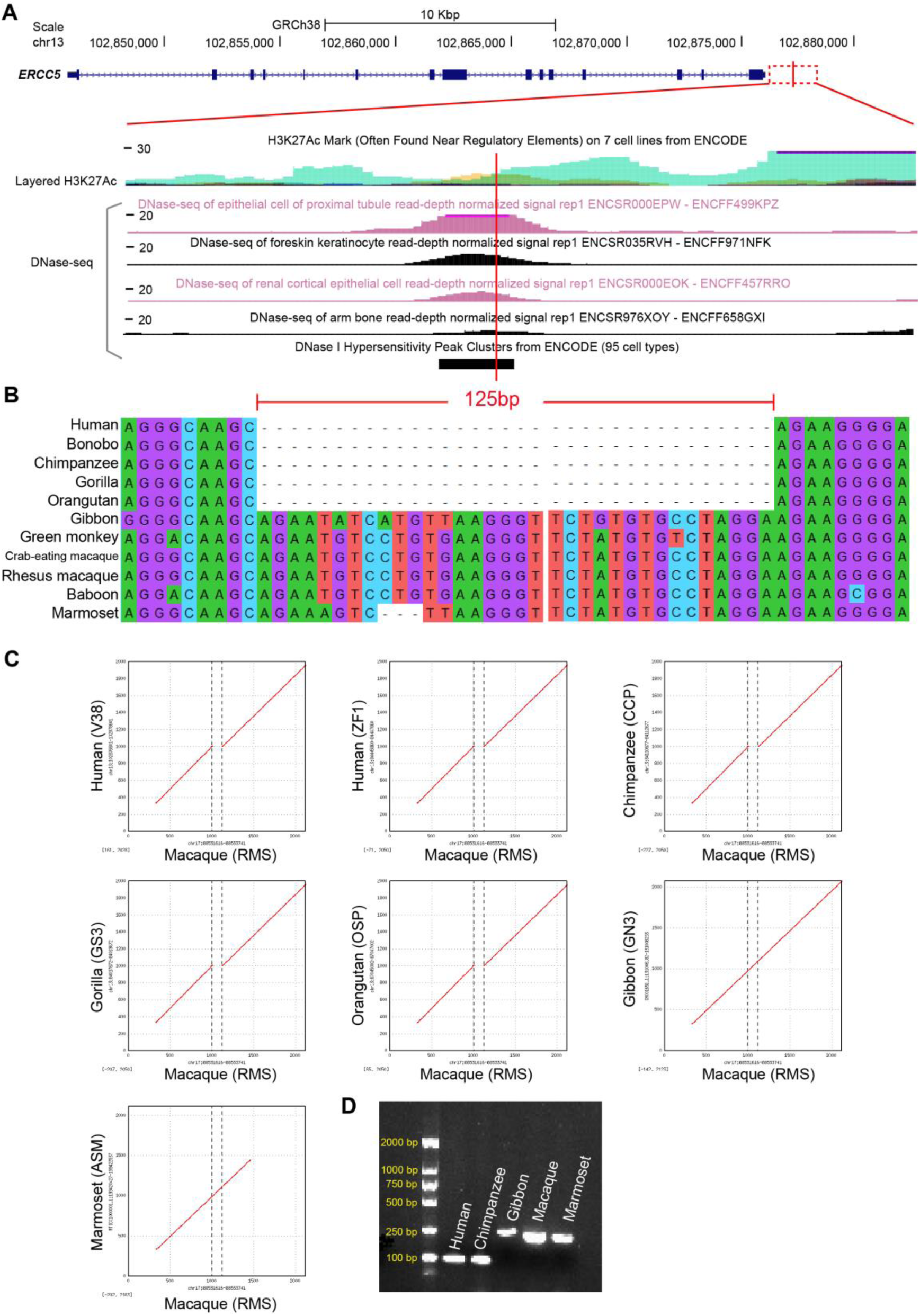
A GASSV (125 bp deletion) located in the regulatory region of *ERCC5*, a gene related to body size. **(A)** Location of the 125 bp deletion in great apes and regulatory annotations (from ENCODE). **(B)** Sequence alignments among primates. **(C)** Dot plot for the pairwise comparison of the deletion region (1 Kbp downstream and upstream flanking sequences) between macaque and apes. **(C)** PCR validation of the 125 bp deletion in *ERCC5*.

**Figure S23.**
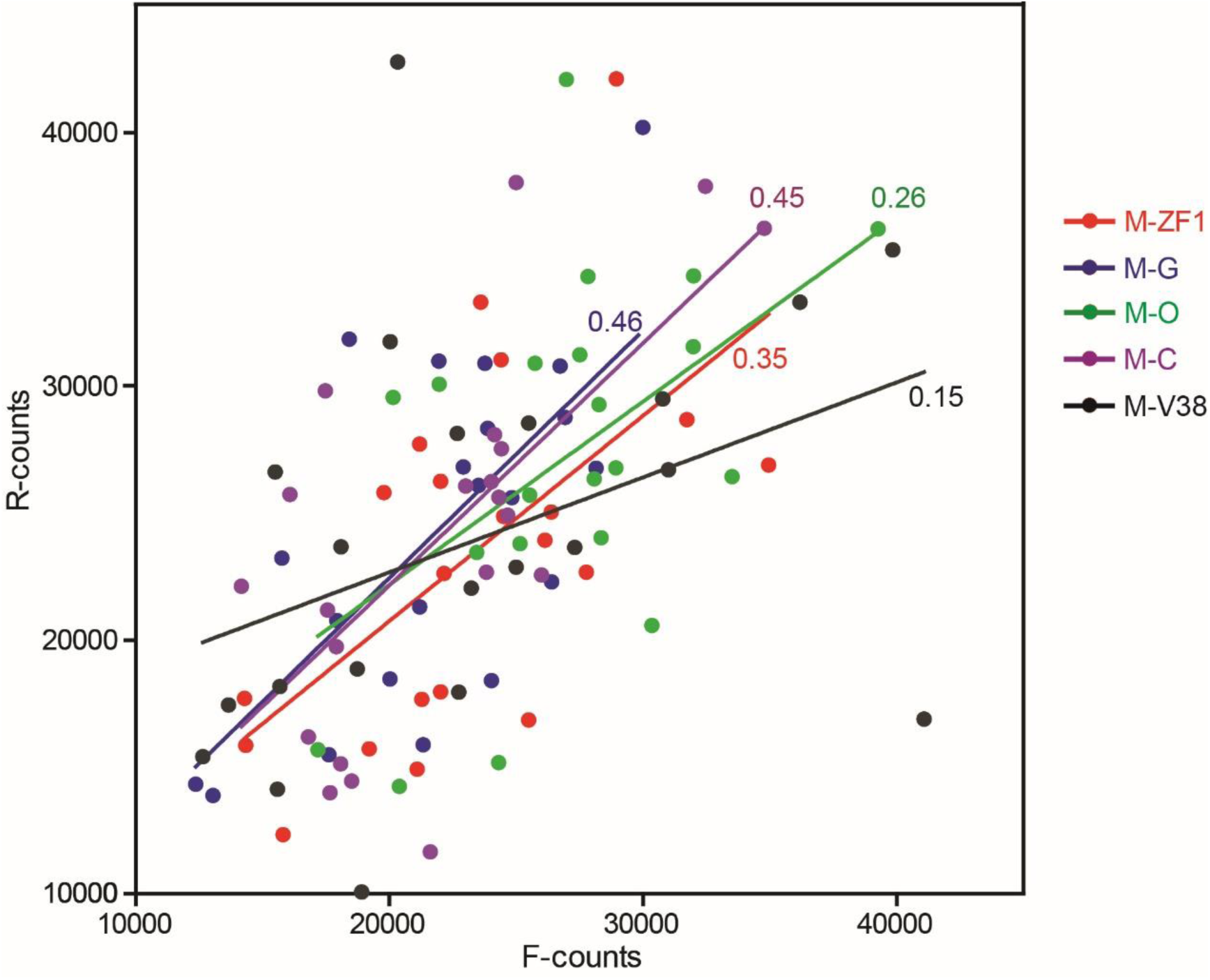
Correlation of the SV numbers between forward-calling and reverse-calling. Genome-comparison-based SV callings are performed (Methods) between macaque (M) and apes, including ZF1-human (a long-read assembly of human), G-gorilla (GS3), O-Orangutan (OSP), C-chimpanzee (CCP) and V38-human (the human reference genome GRCh38). The SV counts are marked in different colors. Each dot refers to the different chromosome. The regression coefficients (R^2^) are indicated.

**Figure S24.**
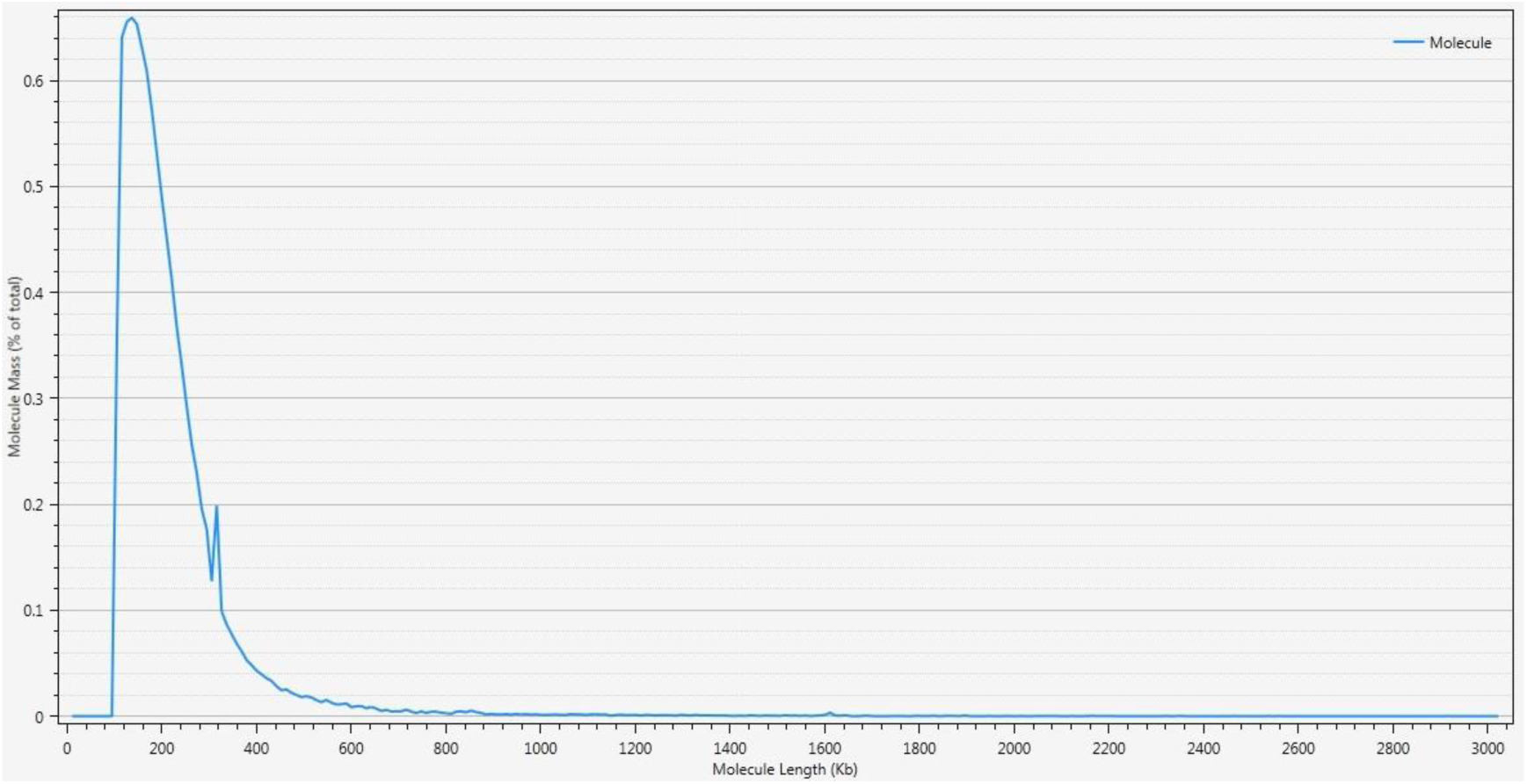
Distribution of molecular length for the Bionano clean data.

**Figure S25.**
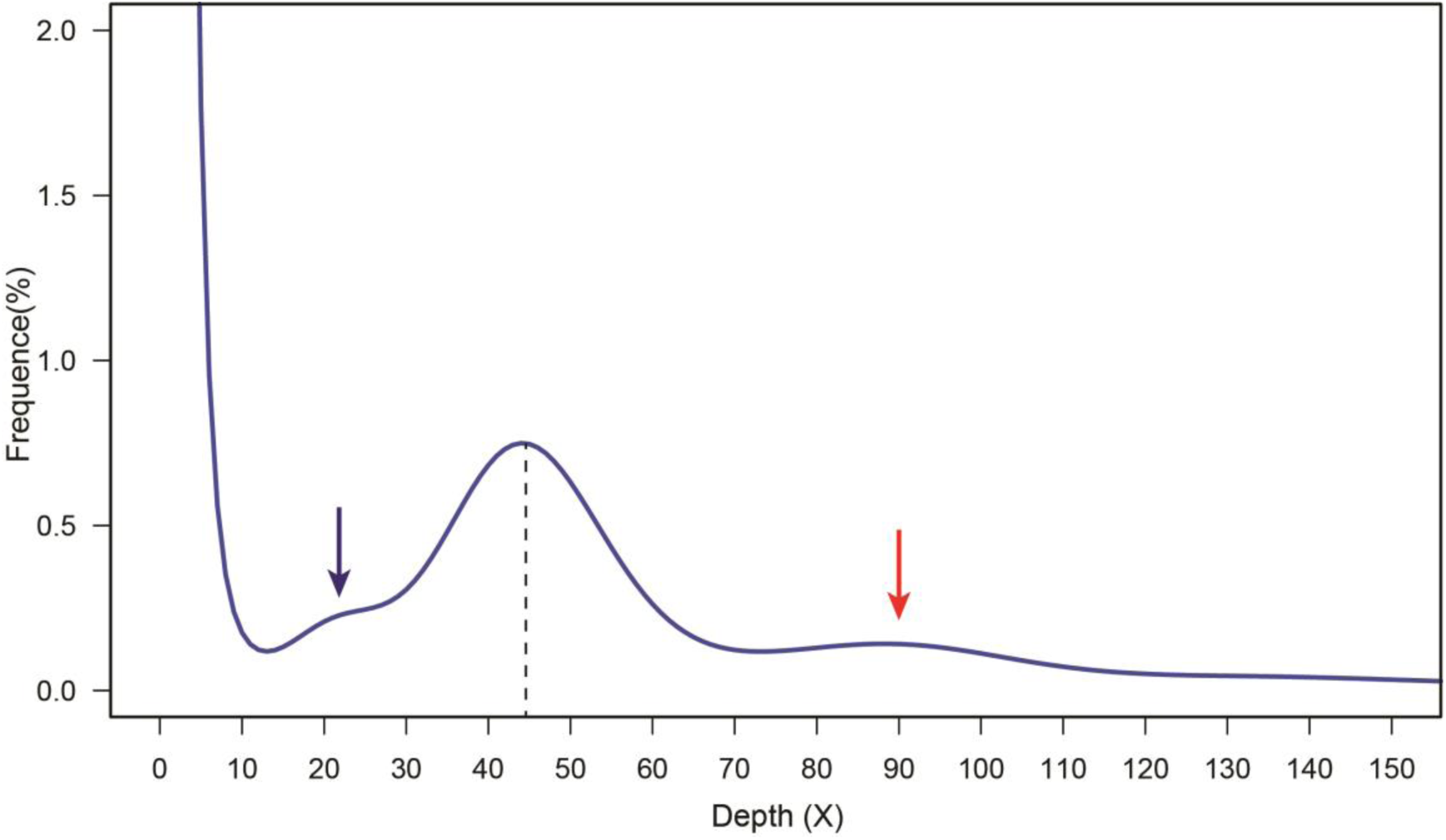
The 17-mer depth distribution curve. K-mer depth is 45× (main peak: MP, vertical dashed line). Two secondary peaks (arrows) at X=1/2*MP and 2*MP coordinates, respectively, suggesting that rheMacS possess a high heterozygosity (arrow in blue) and high repeats (arrow in red).

**Figure S26.**
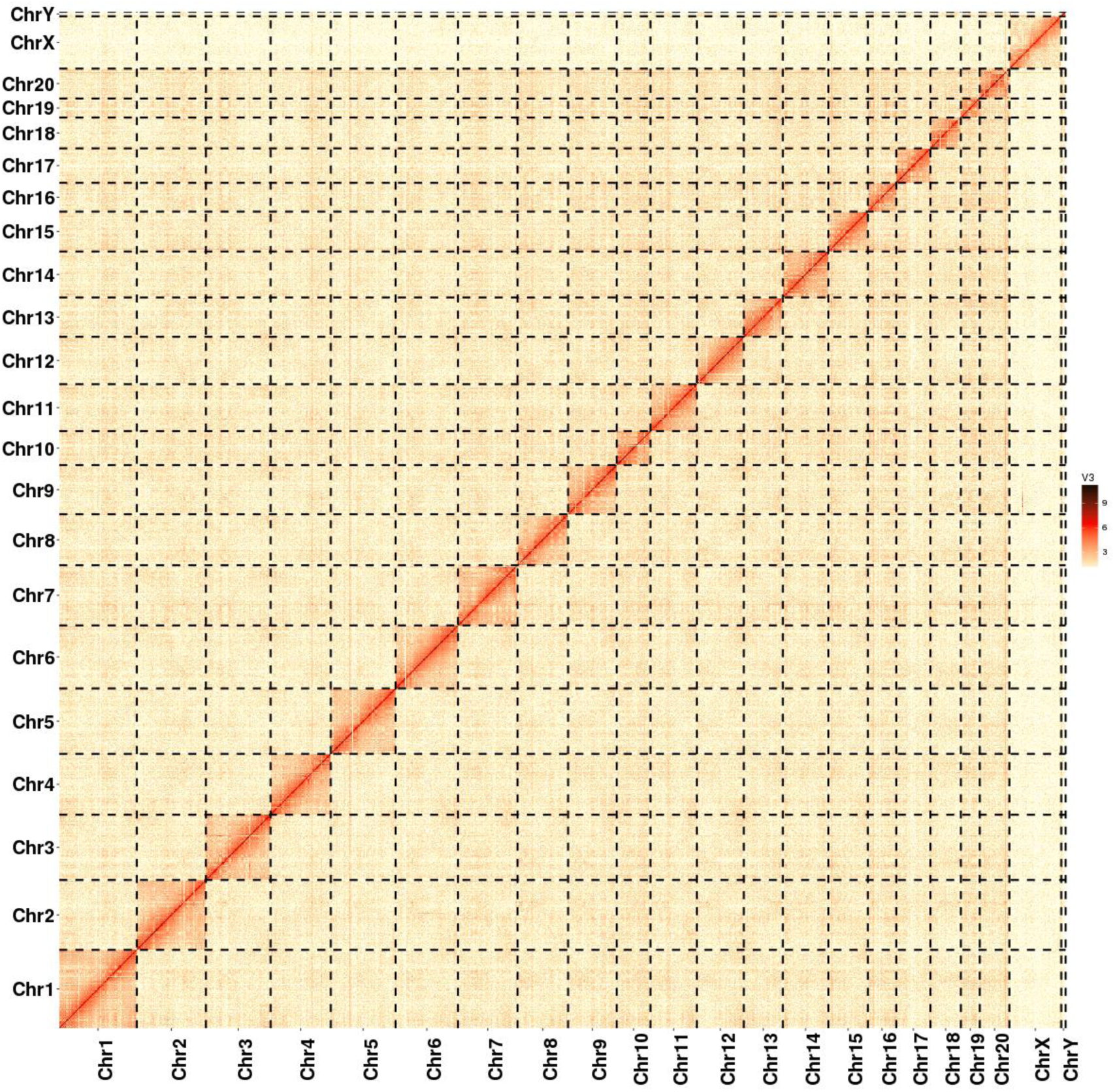
Genome-wide all-by-all chromosome heatmap of the Hi-C data aligned to the rheMacS chromosomes.

**Figure S27.**
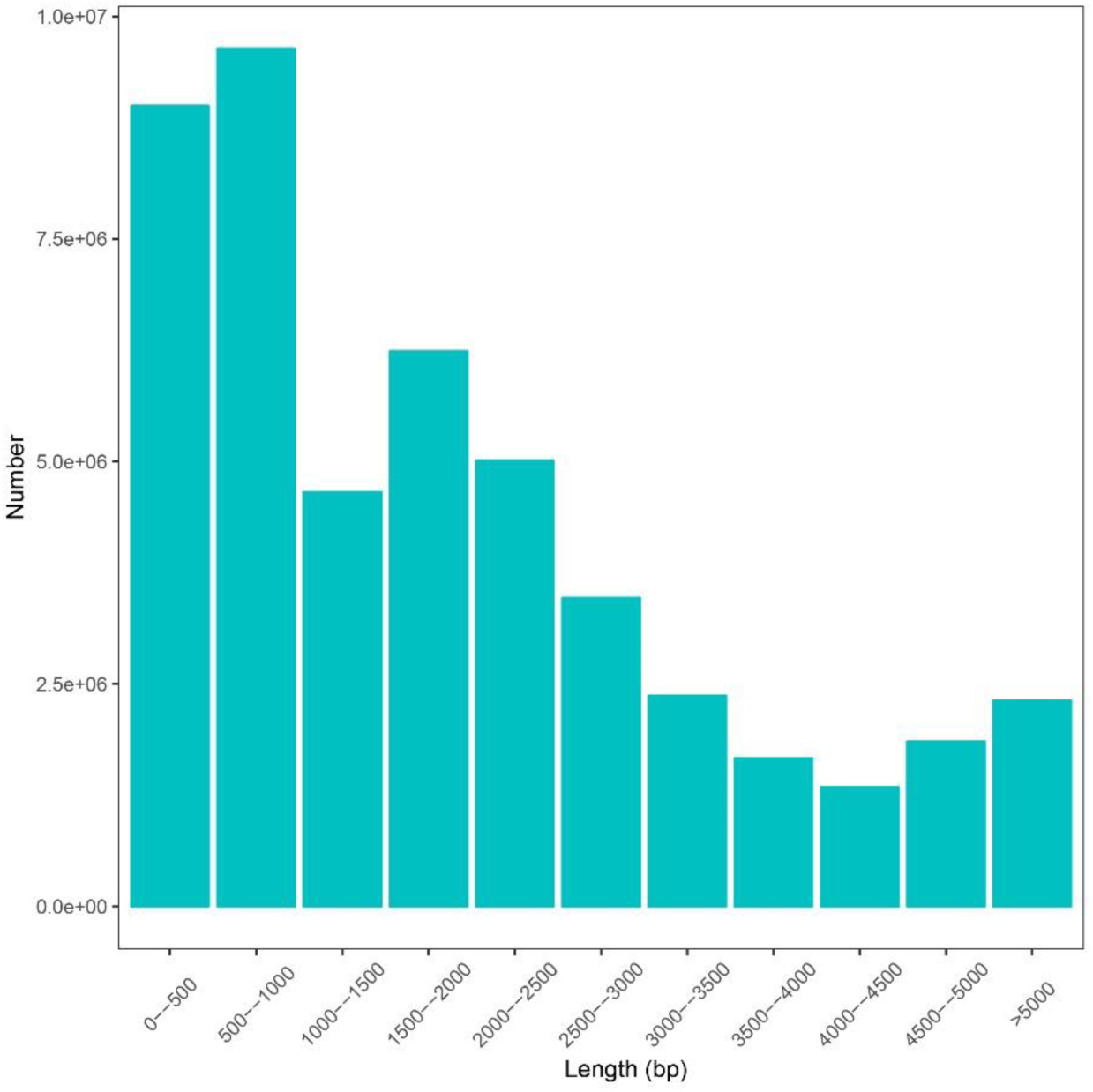
Distribution of the subread lengths of the rheMacS Iso-Seq data.

**Figure S28.**
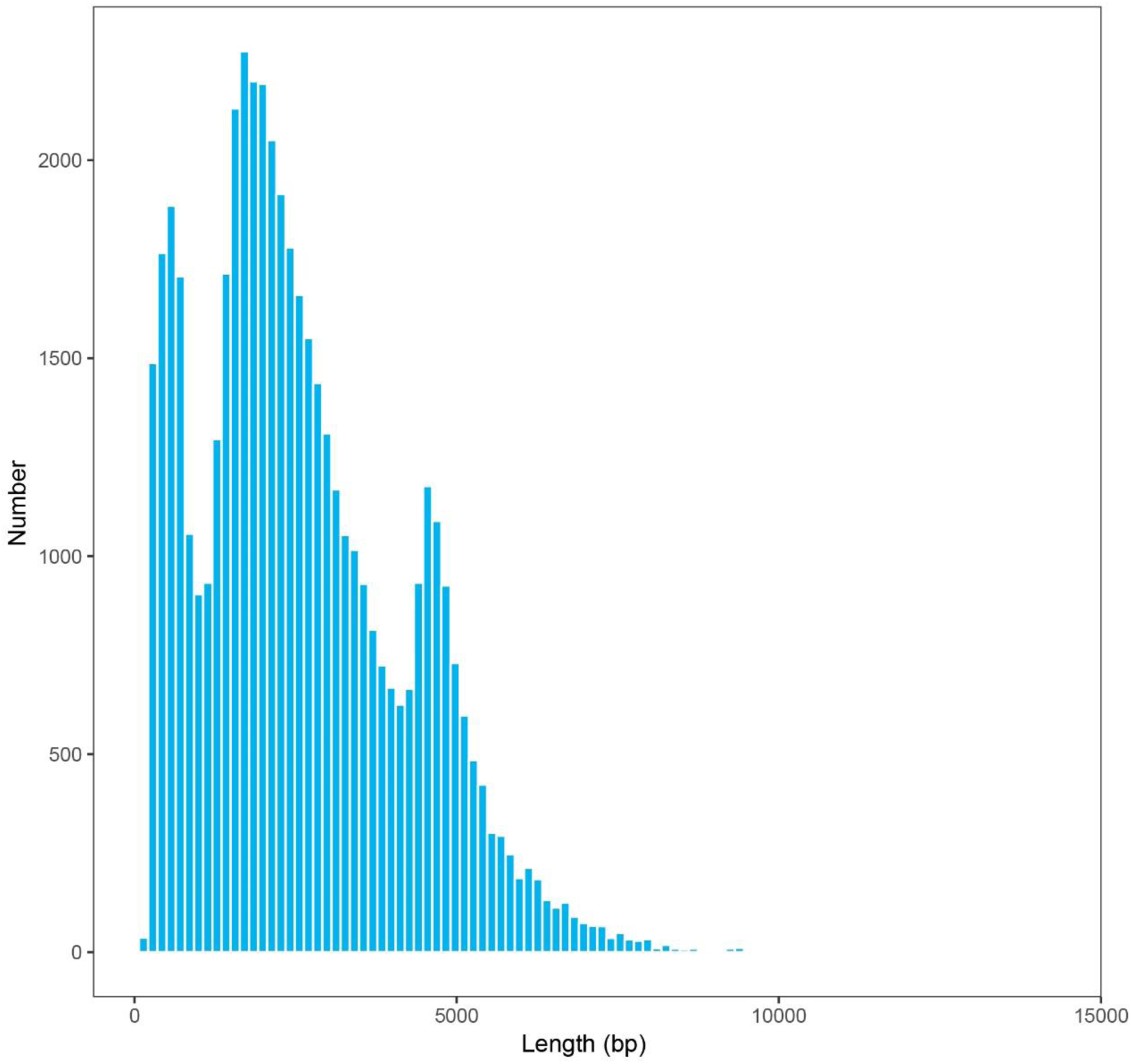
Distribution of the lncRNA length of rheMacS.

